# “Chronic morphine treatment induces a conserved Smchd1-dependent epigenetic memory that disrupts X-chromosome inactivation and genomic imprinting”

**DOI:** 10.64898/2026.03.23.713629

**Authors:** Iraia Muñoa-Hoyos, Manu Araolaza, Irune Calzado, Mikel Albizuri, Paloma Garcia, Nerea Subirán

**Affiliations:** Faculty of Medicine and Nursing, University of the Basque Country (UPV/EHU), 48940. Bizkaia, Spain; Bizkaia Health Research Institute, Barakaldo, 48903. Bizkaia, Spain; Department of Cancer and Genomic Sciences, College of Medicine and Health, University of Birmingham, Birmingham, B15 2TT, UK

**Author notes:** Correspondence: Nerea Subirán Ciudad, Department of Physiology. Faculty of Medicine and Nursing. University of Basque Country. 48940. Leioa, Bizkaia, Spain. Equal contribution (Iraia Muñoa-Hoyos, Manu Araolaza). These authors contributed equally to the work.

**Keywords:** morphine, epigenetic memory, X-chromosome inactivation, genomic imprinting

## Abstract

Epigenetic memory ensures stable inheritance of gene expression patterns critical for embryonic development. Environmental exposures can disrupt this memory, yet the mechanisms remain unclear. Here we demonstrate that chronic morphine exposure induces a persistent transcriptomic and epigenetic memory by repressing *Smchd1*, a key chromatin regulator, in mouse embryonic stem cells, preimplantation embryos, and human induced pluripotent stem cells. This repression compromises maintenance of X-chromosome inactivation and genomic imprinting, leading to sustained dysregulation of developmentally important gene clusters. Morphine-induced epigenetic alterations also involve changes in DNA methylation and histone modifications along the X chromosome and notably increased H3K27me3 at the *`Smchd1`* locus. These findings reveal a conserved mechanism by which opioid exposure disrupts higher-order chromatin architecture and epigenetic memory during early development, potentially contributing to long-term developmental and clinical outcomes.

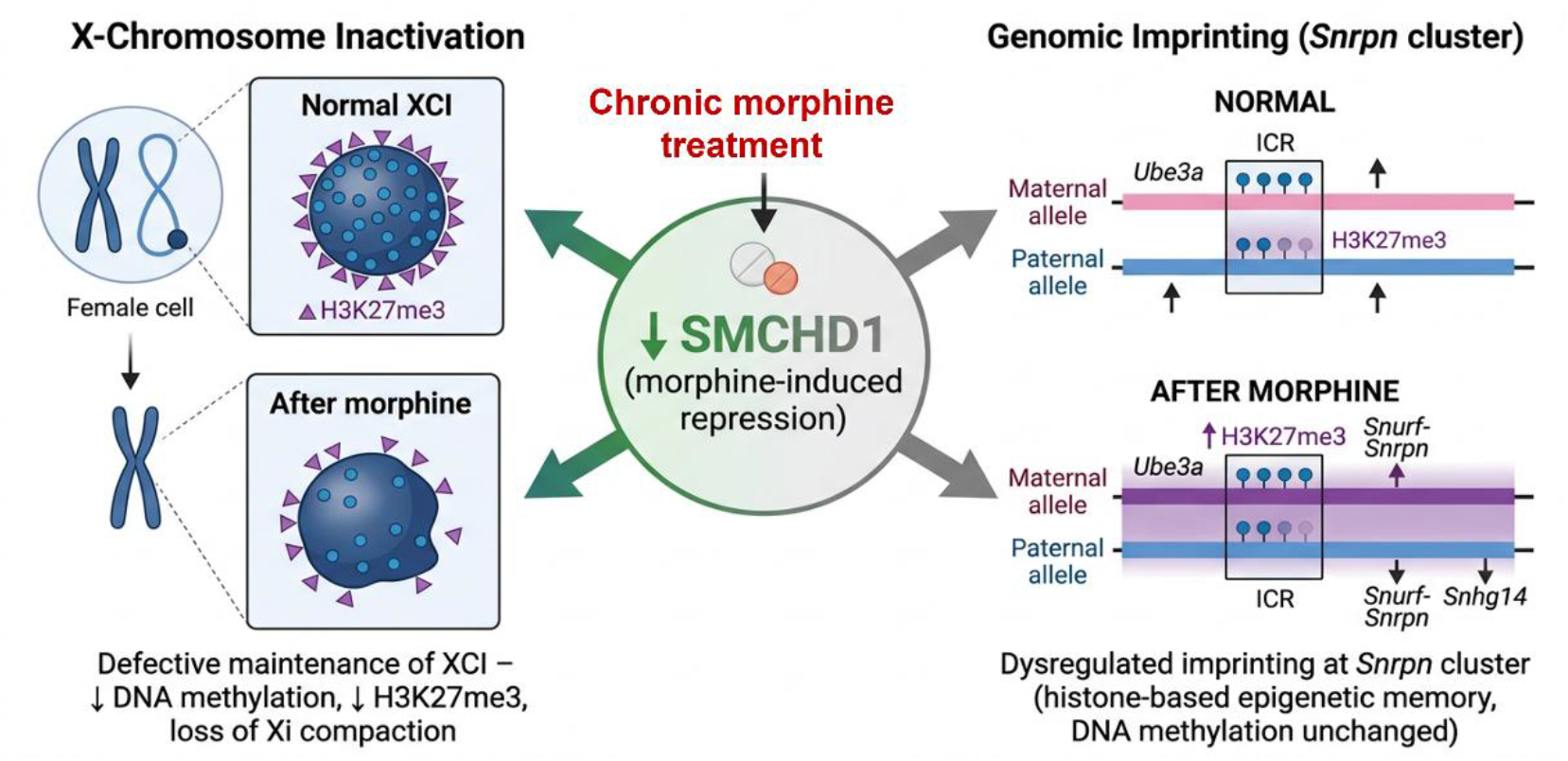

## 1. INTRODUCCION

Cellular memory refers to the intrinsic ability of cells to retain and propagate cell type–specific gene expression patterns across successive cell divisions, even in the absence of the original differentiation cues^1,2^. This phenomenon is fundamental for maintaining cell identity and supporting complex differentiation programs initiated during embryogenesis and sustained throughout life^3^. Specifically, cellular memory enables the formation of tissues and organs from zygote-derived cells that share an identical genome. A key mechanism underlying cellular memory is epigenetic memory, which involves heritable changes in gene expression that persist through mitosis without altering the underlying DNA sequence^4,5^. These epigenetic modifications include DNA methylation, post-translational histone modifications, chromatin remodeling, and the action of non-coding RNAs^2,6–10^. Although these mechanisms are known to play crucial roles in maintaining cell identity, the exact processes by which they ensure the stable inheritance of cellular phenotypes remain incompletely understood. Notably, two processes—X chromosome inactivation (XCI)^11,12^ and genomic imprinting^13^— are widely recognized as classic examples of robust epigenetic memory. Both are established during early development and are stably maintained throughout the organism’s lifespan, ensuring normal development and tissue homeostasis.

Epigenetic memory, therefore, represents a pivotal determinant of physiological development and plays a central role in the etiology of pathophysiological conditions^14,15^. Errors in the formation or maintenance of epigenetic memory or stable epigenetic alterations induced by environmental agents during embryogenesis, can result in aberrant gene activation or silencing, disrupting developmental trajectories and increasing disease susceptibility^16^. Notably, cancer cells can exploit epigenetic memory mechanisms to preserve malignant phenotypes across generations, evading therapeutic interventions^17–19^. Embryonic development is particularly sensitive to environmental exposures, including xenobiotics, pharmacological agents, and or endocrine disruptors among^20,21^, especially those capable of crossing the placental barrier. Among them morphine can cross the placental barrier and reach the developing embryo, inducing anatomical and functional abnormalities, developmental delays, and neurological impairments in rodent models^22,23^. Opioids such as morphine are increasingly recognized for their ability to modulate gene expression, transcription factor activity, DNA methylation, and chromatin architecture, including histone acetylation and methylation, across diverse cell types^24–29^. Notably, opioid exposure during pregnancy has been linked to neurodevelopmental alterations in offspring^22,23^. Despite its widespread use, the molecular mechanisms by which morphine disrupts epigenetic regulation during early development remain poorly understood. Against this backdrop, our study aims to determine whether morphine exposure induces cellular epigenetic memory in mouse embryonic stem cells (mESCs) and to investigate whether this memory persists during lineage commitment and differentiation, thereby providing mechanistic insight into opioid-mediated developmental risks.

## 2. RESULTS

### 2.1. Morphine induces long-term transcriptional reprogramming through sustained epigenetic regulation in mESC

To investigate whether morphine exposure can elicit a persistent epigenetic memory at the cellular level, we first established a tractable *in vitro* model system (Figure 1A). mESCs were selected due to their capacity for indefinite self-renewal and pluripotent differentiation into all somatic lineages, making them an ideal platform to study the impact of environmental stimuli on developmental trajectories. Our aim was to mimic an *in-vitro* chronic morphine treatment, as, it is more likely to produce stable epigenetic changes rather than an acute morphine treatment. As has been previously described^30^, OCT4-reporter mESCs were treated with 10 μM morphine for 24 hours, for chronic treatment, to assess morphine-induced epigenetic modifications (P1). To determine the persistence of these alterations across cellular generations, both treated and untreated cells were reseeded and cultured for an additional 48 hours (P2) and 96 hours (P3) following morphine withdrawal. These time points were selected in accordance with standard mESC passaging intervals, which are required every two days to maintain pluripotency and prevent spontaneous differentiation. No morphological changes nor cell viability were observed at any of the examined time points, as it has been also reported before^31^.

**Figure 1.**
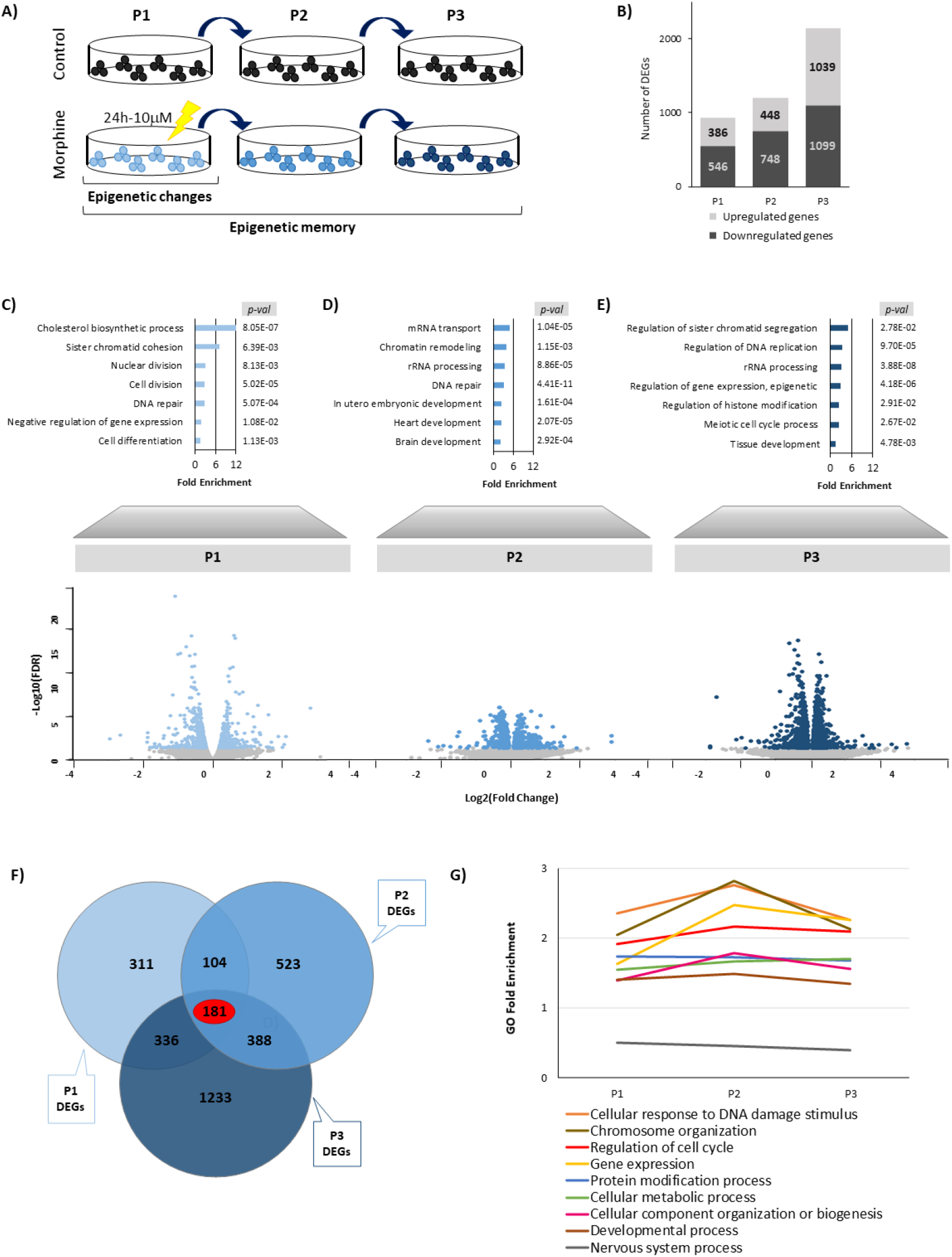
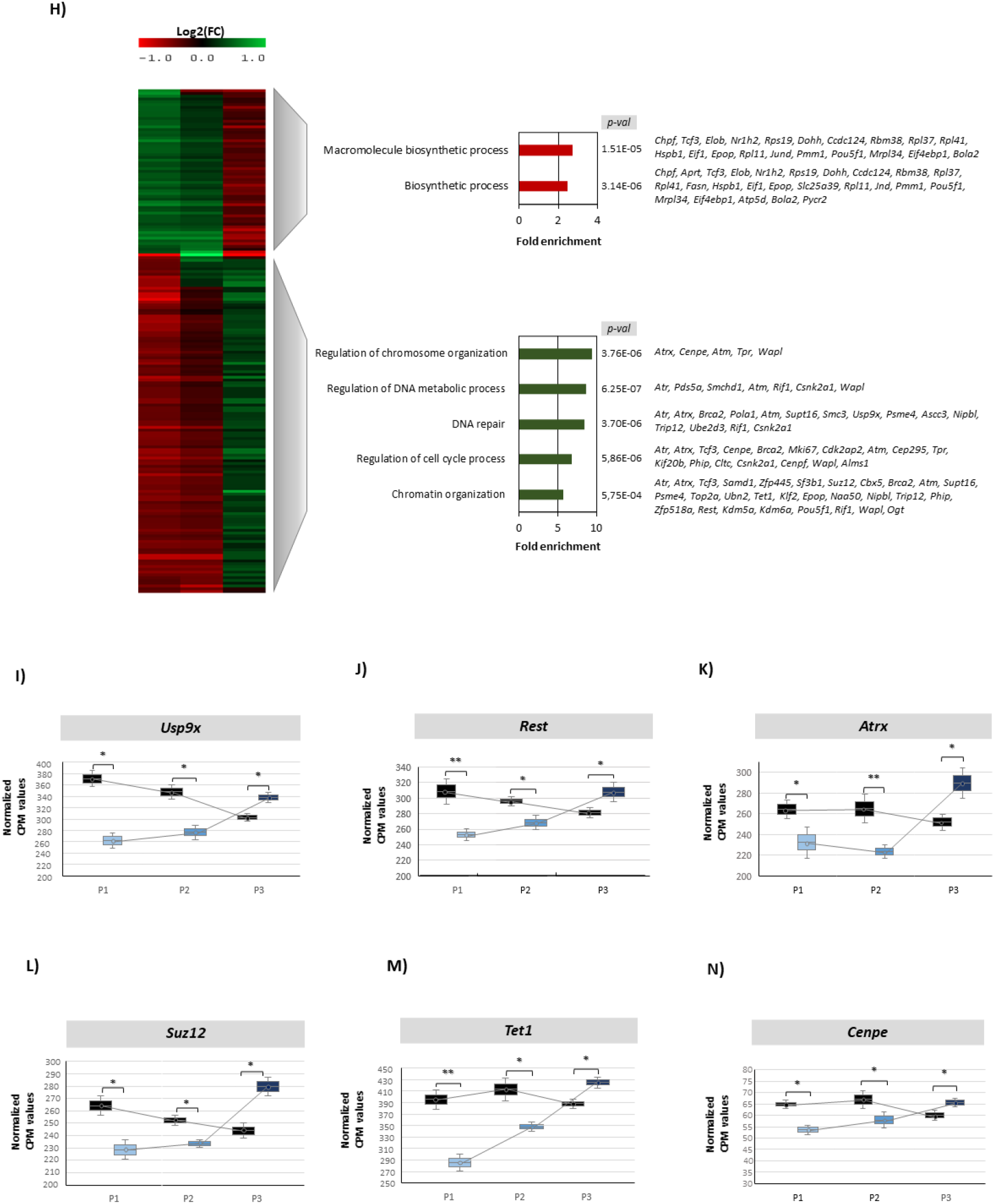
Transcriptomic dynamic after chronic morphine treatment. **(A)** Schematic overview of the experimental design to study the cell-to cell epigenetic memory induced by morphine. mESC collected immediately after chronic morphine treatment (P1; 10 μM, 24 h) and maintained in culture without morphine for additional 48h (P2) and 96h (P3). **(B)** Bar graph showing the number of differentially expressed genes (DEGs; p < 0.05, FDR < 0.05) identified at each passage, including upregulated and downregulated genes. Gene ontology (GO) fold enrichment analyses and volcano plots of DEGs identified (p < 0.05, FDR < 0.05) at **(C)** P1, **(D)** P2, and **(E)** P3. **(F)** Venn diagram illustrating the overlap of DEGs across P1, P2, and P3, highlighting genes detected at multiple time points. **(G)** GO fold enrichment analysis of persistent DEGs, showing enrichment of biological processes across time points. **(H, left)** Hierarchical clustering visualization with heatmap representation of Log2 (FC) values of persistent DEGs shared across passages. **(H, right)** GO enrichment analysis grouped into two clusters according to expression patterns over time. Dynamic expression of **(I)** *Usp9x*, **(J)** *Rest*, **(K)** *Atrx*, **(L)** *Suz12*, **(M)** *Tet1*, and **(N)** *Cenpe* genes across passages P1–P3. Statistical significance between control and morphine for each time points was set at *p < 0.05, **p < 0.01, ***p < 0.001.

We first performed a dynamic transcriptomic analysis of mESCs at the three time points, to characterize morphine-induced transcriptional changes indicative of cellular memory. Principal component analysis (PCA) of global gene expression profiles revealed a clear separation between morphine-treated and control cells across all time points (Supplementary Data 1A). Read count distribution plots confirmed appropriate TMM normalization across samples (Supplementary Data 1B). Differential gene expression analysis identified a progressive increase in the number of differentially expressed genes (DEGs) over time—932 at P1, 1,196 at P2, and 2,138 at P3— suggesting sustained transcriptomic reprogramming following morphine withdrawal (Figure 1B). Applying stringent thresholds (p < 0.05, FDR < 0.05), we found that at P1, 41% of DEGs were upregulated (386 genes) and 59% were downregulated (546 genes), with enrichment in core cellular functions including cell division and differentiation, nuclear division, gene expression, and cholesterol biosynthesis (Figure 1C). After morphine withdrawal, at P2, 37% of DEGs were upregulated (448 genes) and 63% were downregulated (748 genes), mainly associated with chromatin remodeling, RNA processing, DNA repair, and embryo development (Figure 1D). By P3, 49% of DEGs were upregulated (1,039 genes) and 51% were downregulated (1,099 genes), predominantly linked to chromatid segregation and meiosis, DNA replication, epigenetic regulation, among other processes (Figure 1E). These findings suggest that cells retain a memory of prior morphine exposure and undergo transcriptional reprogramming even in the absence of the drug in the culture medium. Indeed, morphine elicits long-lasting changes in the expression of genes encoding epigenetic and meiotic cell cycle regulators, implying a deregulation of epigenetic mechanisms that may underlie cellular memory.

To further elucidate the persistence of morphine-induced alterations in these functions, we performed integrative transcriptomic analyses across three time points. Venn diagram analysis identified 181 DEGs consistently deregulated throughout the time course (Figure 1F). Gene ontology enrichment revealed that nuclear-related functions, including DNA damage response, chromosome organization, gene expression regulation, and cell cycle control, exhibited the highest fold enrichment and remained persistently affected following morphine withdrawal (Figure 1G). In addition, key cellular processes such as protein modification, metabolism, and biogenesis were also disrupted. Notably, although genes associated with development and nervous system function showed lower enrichment scores, their expression remained consistently altered across all time points (Figure 1G). This pattern suggests that morphine exposure induces a sustained transcriptomic signature, indicative of epigenetic memory, with a primary impact on neuronal developmental gene networks. Hierarchical clustering of persistent DEGs revealed distinct transcriptional profiles between morphine-treated and control mESCs, classifying the persistent DEGs into two major clusters (Figure 1H, Supplementary Data 2). The first cluster comprised genes primarily involved in macromolecule biosynthetic processes, which were upregulated immediately following morphine exposure but progressively downregulated by P3. In contrast, the second cluster included genes that were initially downregulated and gradually increased in expression, reaching upregulation at P3 (Figure 1H, Supplementary Data 2). Ontology analysis indicated that this cluster was enriched in nuclear function such as DNA metabolism and repair, cell cycle process and chromosome and chromatin organization. Among the most significantly affected regulators in this cluster were key cell fate determinants involved in neuronal development, such as *Usp9x, Rest* and *Atrx* (Figure 1I-K), as well as several epigenetic regulators including *Suz12* and *Tet1* (Figure 1L and M) and important cell cycle regulators involved in mitotic progression such as *Cenpe* (Figure 1N). These transcriptional dynamics suggest a biphasic response to morphine, indicative of transcriptomic reprogramming consistent with a potential long-term epigenetic cellular memory.

### 2.2. Morphine Induces Sustained Repression of Epigenetic Regulator *Smchd1* in mESCs

To investigate the epigenetic mechanisms underlying the persistent transcriptional alterations observed in morphine-treated mESCs, we first performed an integrative multi-omics analysis combining transcriptomic (RNA-seq), methylome (whole-genome bisulfite sequencing, WGBS), and the repressive histone H3K27me3 (ChIP-seq) data (Figure 2A). Specifically, the 181 persistent DEGs identified across the time course following morphine exposure were compared with 15,357 morphine-induced differentially methylated cytosines (DMCs) and 1,024 differentially enriched H3K27me3 regions (DBSs). Venn diagram revealed 14 genes significantly affected by morphine at both the transcriptomic and epigenetic levels, eight of which encode key transcriptional end epigenetic regulators (Figure 2B, Supplementary Data 3). Notably, *Smchd1*, a gene implicated in genome-wide epigenetic repression and essential for maintaining silencing of the inactive X chromosome (Xi) in female mammals^32^, emerged as a key target of morphine-induced epigenetic disruption. Genome browser visualization of the *Smchd1* locus confirmed alterations at both transcriptomic and epigenetic levels between control and morphine-treated mESCs (Figure 2C). RNA-seq analysis demonstrated a marked reduction in *Smchd1* transcript levels following morphine exposure (Figure 2C and 2D). Although we did not find any methylation changes at promotor region, WGBS revealed a global hypomethylation across the gene body, which also involved a CTCF region (Figure 2C and 2E). In contrast, morphine induced a hypermethylation in an enhancer region downstream of the TSS, potentially contributing to transcriptional silencing (Figure 2C and 2E). Additionally, ChIP-seq analysis revealed significant enrichment of the repressive histone mark H3K27me3 not only at the *Smchd1* promoter but also in the same enhancer region in morphine-treated cells (Figure 2C and 2F). These findings indicate that morphine downregulates *Smchd1* through a combined epigenetic repression, which involves the modulation of methylation levels as well as the increase of H3K27me3 deposition at different regulatory elements of the loci.

**Figure 2.**
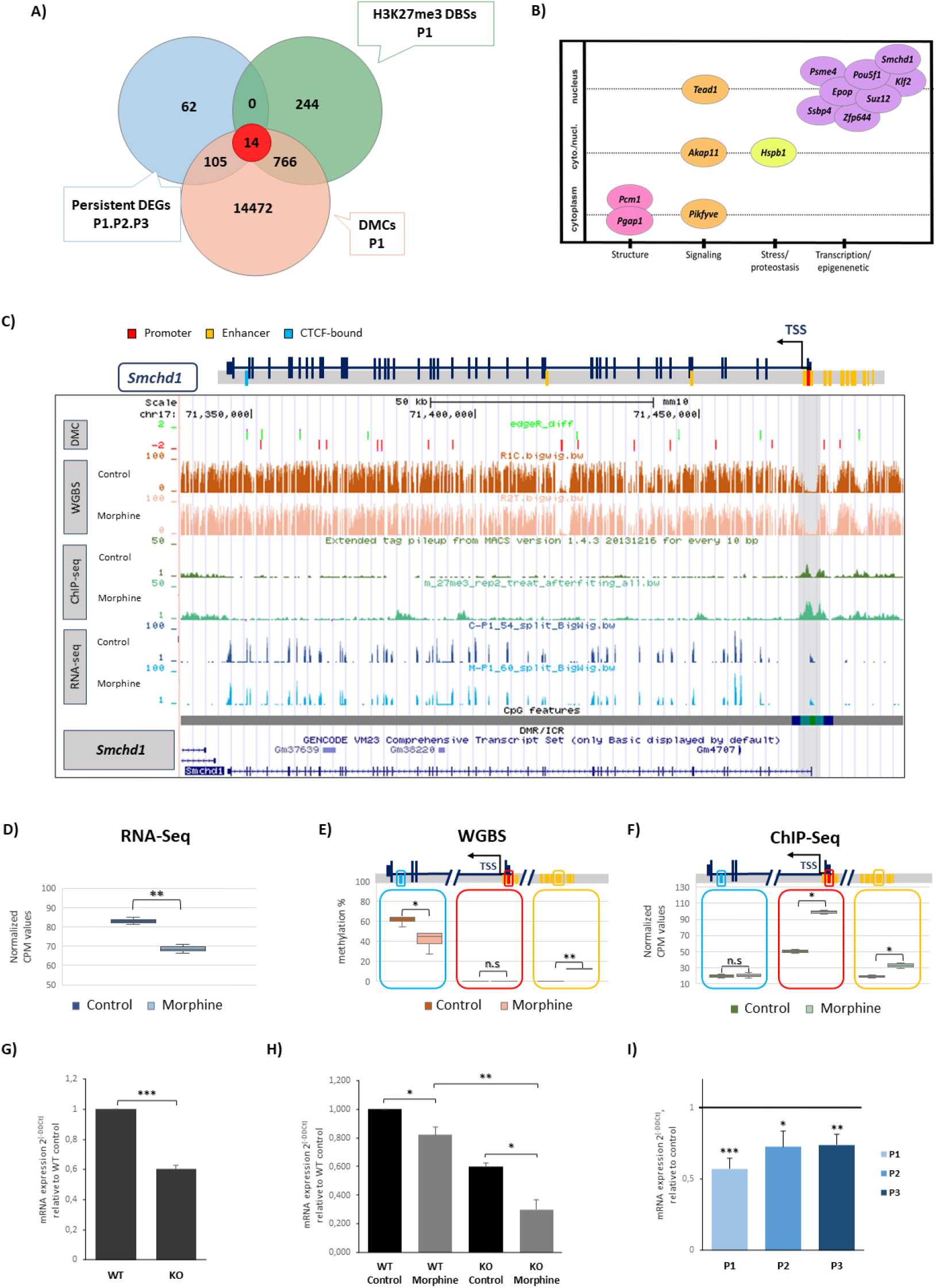
Effect of chronic morphine treatment on Epigenetic Repressor *Smchd1* gene. **(A)** Venn diagram showing the overlap between differentially expressed genes (DEGs) common to passages P1–P3, differentially methylated cytosines (DMCs) identified at P1, and H3K27me3 differential binding sites (DBSs) identified at P1. **(B)** Functional interaction network constructed based on molecular function, biological process, and cellular localization of genes identified across the three high-throughput sequencing approaches. **(C)** Genome browser tracks at the *Smchd1* locus showing WGBS obtained methylation levels, H3K27me3 ChIP-seq signal, and RNA-seq coverage under control and morphine-treated conditions. Regulatory annotations, conserved elements, and CpG features (CGIs, shores, shelves, and open sea regions) are shown below, together with a schematic representation of the gene structure. Box plots represent **(D)** *Smchd1* gene expression as counts per million, **(E)** % of DNA methylation and **(F)** H3K27me3 levels at *Smchd1* CTCF-bound (blue), promoter (red), and enhancer (yellow) regions, for each condition. Bar plots represent RT-qPCR analysis of *Smchd1* gene expression **(G)** in wild-type and partial Smchd1 knockout mESCs, **(H)** in wild-type and partial *Smchd1* knockout mESCs following morphine exposure **(I)** and across passages P1–P3. Statistical significance between control and morphine was set at *p < 0.05, **p < 0.01, ***p < 0.001.

To further validate *Smchd1* as a morphine-sensitive target, we employed a gene-silencing approach (Supplementary Data 4). Using CRISPR/Cas9-mediated gene editing, we generated partial *Smchd1* knockout mESCs (*Smchd1*^*+*^*/*^*−*^) via plasmid-based transfection^33^, enabling assessment of reduced rather than complete gene loss, as full knockout would preclude evaluation of morphine’s effects. RT-qPCR confirmed a partial decrease in *Smchd1* expression, validating the editing strategy (Figure 2G). Upon chronic morphine treatment, *Smchd1*^*+*^*/*^*−*^ cells exhibited a significantly greater reduction in transcript levels compared with wild-type controls, demonstrating that the remaining allele is highly sensitive to morphine (Figure 2H). Finally, dynamic RT-qPCR across consecutive passages (P1–P3) confirmed sustained *Smchd1* downregulation following morphine exposure (Figure 2I). Collectively, these results identify *Smchd1* as a morphine-sensitive chromatin regulator whose prolonged repression may disrupt X-chromosome inactivation dynamics in mESCs, indicating that morphine exposure compromises higher-order chromatin organization and alter epigenetic homeostasis at the long-term.

### 2.3. Morphine Impairs the Maintenance of X Chromosome Inactivation

Owing the critical role of SMCHD1 in maintaining the silencing of the Xi in female mammals ^35,36^, we next sought to evaluate the impact of morphine on X-chromosome inactivation processes. First, we examined the expression and epigenetic regulation of *Xist*, the long non-coding RNA that initiates XCI by spreading in cis along the X-chromosome and recruiting Polycomb repressive complexes PRC1 and PRC2 (Figure 3A). In the initiation phase, *Xist* is up-regulated from the future inactive X, while its antisense transcript *Tsix*, which overlaps the same locus, is down-regulated. This switch marks the onset of chromosome-wide silencing^34,35^. RNA-seq analysis revealed no significant changes in *Xist* or *Tsix*, placed in the same loci, following morphine exposure (Figure 3B–D). Consistently, DNA methylation patterns and the distribution of the repressive histone mark H3K27me3 across the locus encompassing both genes remained unaffected after chronic morphine treatment (Figure 3B). Next, we evaluated the levels of repressive histones H2AK119ub and H3K27me3, which are catalyzed by PRC1 and PCR2 respectively during the initiation of XCI. Although H2AK119ub levels were not significantly altered, we observed a marked reduction in H3K27me3 in morphine-treatment cells (Figure 3E). To validate this impact on XCI process, we analyzed the distribution of this repressive histone specifically along the X-chromosome. The X-chromosome landscape revealed also a global decrease in H3K27me3 enrichment along the chromosome after morphine treatment (Figure 3F and 3G), consistent with impaired PRC2 activity (Figure 3H). In fact, RT-qPCR analysis showed down-regulation of *Ezh2*, the catalytic subunit of PRC2, after 24 h of morphine exposure (P1), but returned to control levels by 96 h (P3) (Figure 5C). These results indicate that morphine disrupts the epigenetic landscape associated with the initiation of X-chromosome inactivation, primarily by reducing PRC2 activity and H3K27me3 deposition of the chromosome, without directly altering *Xist* or *Tsix* expression.

**Figure 3.**
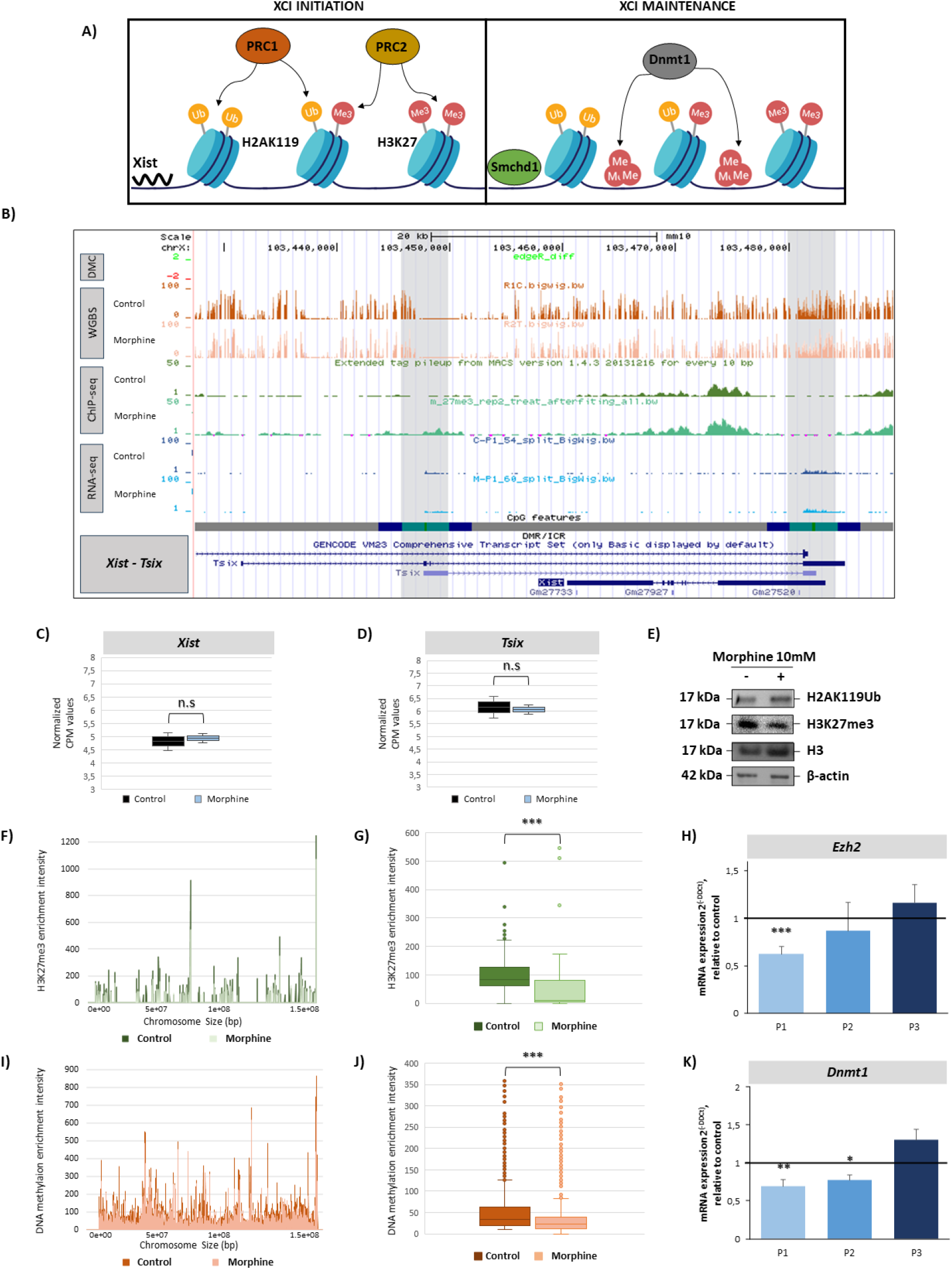
Effect of chronic morphine treatment on X-chromosome inactivation process. **(A)** Schematic overview of X-chromosome inactivation (XCI) process, illustrating initiation and maintenance phases and the associated chromatin regulators. **(B)** Genome browser tracks of the *Xist/Tsix* locus showing WGBS obtained methylation levels, H3K27me3 ChIP-seq signal, and RNA-seq coverage under control and morphine-treated conditions. Regulatory annotations, conserved elements, and CpG features (CGIs, shores, shelves, and open sea regions) are shown below, together with a schematic representation of the genes structure. Box plots represent **(C)** *Xist* and **(D)** *Tsix* gene expression as counts per million. **(E)** Western blot analysis of H2AK119ub and H3K27me3 protein levels in control and morphine-treated mESCs. H3 and β- actin is used as a loading control. **(F)** Landscape of repressive histone H3K27me3 along the X chromosome in control and morphine-treated mESCs. **(G)** H3K27me3 ChIP-seq enrichment of the X chromosome after chronic morphine treatment. **(H)** Bar plots represent RT-qPCR analysis of *Ezh2* expression across passages P1–P3. **(I)** Landscape of methylation levels along the X chromosome in control and morphine-treated mESCs. **(J)** DNA methylation levels of the X chromosome after chronic morphine treatment. **(K)** Bar plots represent RT-qPCR analysis of *Dnmt1* expression across passages P1–P3. Statistical significance between control and morphine was set at *p < 0.05, **p < 0.01, ***p < 0.001.

During maintaining phase of the XCI, DNA methyltransferase DNMT1 and the SMCHD1 cooperate to preserve the silenced chromatin state through DNA methylation and higher-order chromatin compaction (Figure 3A). These coordinated epigenetic mechanisms ensure the stability of the inactive X-chromosome (Xi) and its faithful propagation across successive cell divisions. Given that morphine down-regulated *Smchd1* expression—a critical factor for XCI maintenance—we next assessed the impact of chronic morphine treatment on DNA methylation. WGBS also revealed a global reduction in DNA methylation across the X-chromosome (Figure 3I and 3J). RT-qPCR analysis further confirmed that *Dnmt1* expression was down-regulated following morphine exposure and remained suppressed for at least 48 h (P2) but not at 96 h (P3) after morphine withdrawal (Figure 3K). These findings indicate that morphine compromises the maintenance of X-linked gene silencing not only by reducing Smchd1 expression but also through DNA hypomethylation of the X-chromosome.

### 2.4. Morphine Induces Dysregulation of Imprinted Gene Clusters with a Persistent effect at the *Snrpn* Locus

As imprinting is another key process of epigenetic memory during embryonic development, in which certain gene clusters are known to be repressed by SMCHD1^36^, we next investigated whether chronic morphine exposure affects genomic imprinting. RNA-seq data focusing exclusively on imprinted genes confirmed that chronic morphine exposure induced time-dependent and persistent alterations in imprinted gene expression (Figure 4). At 24 h (P1), only four imprinted genes were differentially expressed, increasing to 12 and 15 at P2 and P3, respectively after morphine removal (Figure 4A). At P1, differentially expressed genes were predominantly maternally expressed, whereas paternally expressed genes became more prevalent at P2, although maternally expressed genes exhibited the largest fold changes. By P3, all three imprinting classes were represented, with generally smaller effect sizes (Figure 4B), reflecting a dynamic yet temporally ordered imprinting response, even in the absence of morphine in the medium. Analysis of DEGs across time points revealed no single gene consistently affected at all stages (Supplementary Data 5A). However, cluster-level analysis confirmed an overlap across the three time points. Persistent deregulation across multiple imprinted clusters were highlighted, including *Dlk1–Dio3, Kcnq1, Snrpn, Peg3, Sgce–Peg10, Gpr1–Zdbf1, H13–Mcts2, Igf2–H19, Igf2r–Air, Peg1–Copg2*, and *Tsix–Zcchc13* (Supplementary Data 5B). This underscores a robust role in maintaining morphine-induced epigenetic memory through imprinting mechanisms.

**Figure 4.**
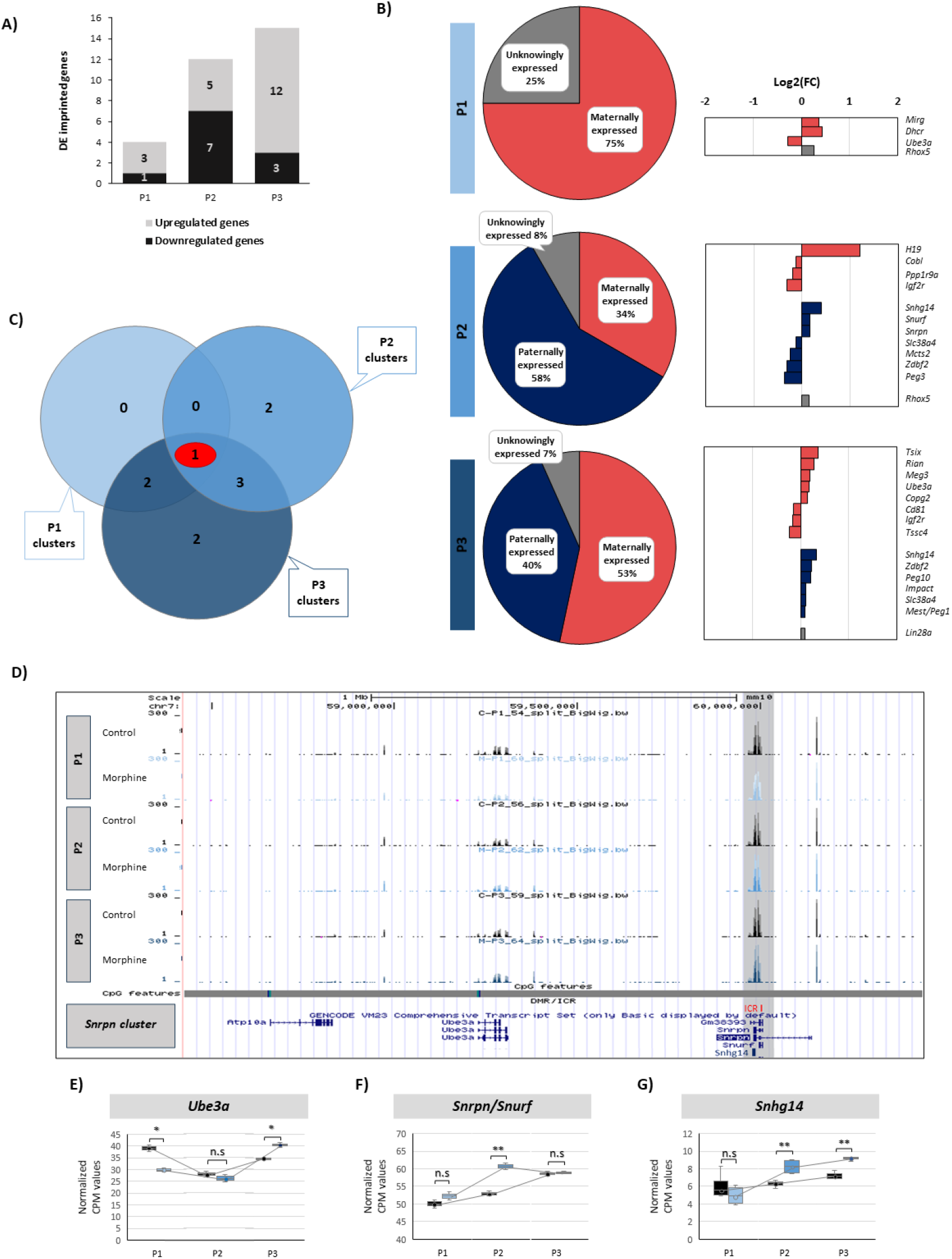
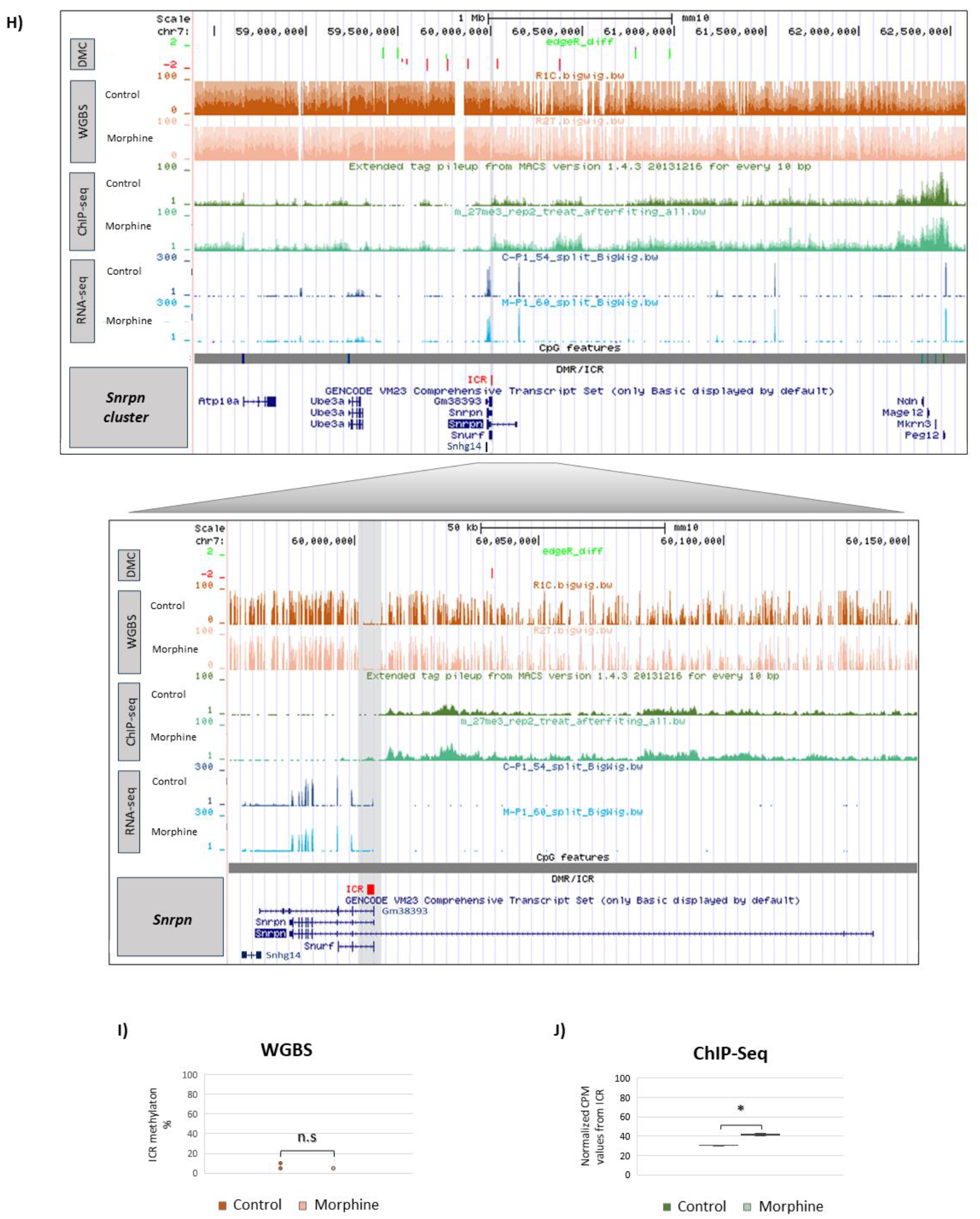
Effect of chronic morphine treatment on Imprinted Gene Clusters. **(A)** Number of differentially expressed imprinted genes identified at passages P1–P3. **(B)** Distribution of maternally expressed, paternally expressed, and biallelically expressed imprinted genes at each passage, together with log_2_ fold-change values (p < 0.05, FDR < 0.05). **(C)** Venn diagram showing overlap of deregulated imprinted clusters across passages. **(D)** Landscape of the *Snrpn* imprinted cluster showing RNA-seq profiles at passages P1–P3. Box plots represent **(E)** *Ube3a*, **(F)** *Snrpn-Snurf*, and **(G)** *Snhg14* gene expression as counts per million across passages (P1-P3). **(H)** Genome browser tracks at the *Snrpn* cluster showing WGBS obtained methylation levels, H3K27me3 ChIP-seq signal, and RNA-seq coverage under control and morphine-treated conditions. Regulatory annotations, conserved elements, and CpG features (CGIs, shores, shelves, and open sea regions) are shown below, together with a schematic representation of the gene structure. Box plots represent **(I)** % of DNA methylation and **(J)** H3K27me3 levels at the *Snrpn* imprinting control region (ICR). Statistical significance between control and morphine was set at *p < 0.05, **p < 0.01, ***p < 0.001.

Notably, the *Snrpn* cluster was uniquely affected at all three time points (Figure 4C). Analysis of the *Snrpn* locus revealed that maternally expressed *Ube3a*, paternally expressed *Snurf* and *Snrpn* —both genes placed at the same loci— and the long non-coding RNA *Snhg14* were expressed in mESCs, whereas the maternally expressed *Atp10a* was not detected (Figure 4D). Chronic morphine exposure induced dynamic, gene-specific transcriptional changes among expressed imprinted genes of the *Snrpn* cluster. *Ube3a* exhibited an initial downregulation at P1, followed by increased expression at P3 (Figure 4E). In contrast, the paternally expressed gene *Snurf-Snrpn* were upregulated at P2 (Figure 4F), and the long non-coding *Snhg14* was also upregulate at P2, remaining elevated at P3 (Figure 4G). Although DNA methylation remained unaltered along the *Snrpn* cluster, including at the imprinting control region (ICR) (Figure 4H and 4I), chip-Seq data of the repressive histone mark H3K27me3 revealed that morphine-induced transcriptional perturbations were accompanied by epigenetic remodeling within *Snrpn* cluster (Figure 4H). The H3K27me3 enrichment increased across the entire cluster locus after morphine treatment and, remarkably, morphine significantly increased the H3K27me3 distribution at the *Snurf-Snrpn* gen promoter, which corresponds to the ICR (Figure 4H, 4J). These findings demonstrate that chronic morphine exposure disrupts imprinting regulation in a temporally ordered and cluster-specific manner, with the *Snrpn* locus emerging as a particularly sensitive and persistent target. The uncoupling between stable DNA methylation and increased H3K27me3 enrichment indicates that morphine-induced epigenetic memory at imprinted loci is mediated predominantly through histone-based mechanisms, revealing a previously unappreciated layer of imprinting plasticity in response to opioid exposure during early embryonic stages.

### 2.5. Morphine-Induced Downregulation of *Smchd1* Is Conserved Across Developmental Stages and Species

To determine whether the morphine-induced repression of *Smchd1* observed in mESCs is maintained across developmental transitions and conserved between species, we performed a series of complementary *in vitro* and *ex viv*o assays (Figure 5). As pluripotent stem cells progress toward an epiblast-like state to preserve developmental competence, we aim to determine if morphine induces sustained repression of *Smchd1* during epiblast-like cells differentiation. To address this goal, mESCs were pretreated with morphine (10 μM, 24 h) and then driven *in vitro* toward an epiblast-like cell (mEpiLC) state in the absence of the drug. Morphological assessment confirmed efficient differentiation in both conditions, as cells transitioned from the characteristic domed mESC colony morphology to the flattened and compact appearance typical of mEpiLCs. Morphine treatment did not induce any observable morphological changes in either mESCs or mEpiLCs (Figure 5A). Exit from pluripotency was further validated by the expected induction of *Dnmt3b* (Figure 5B) and concomitant downregulation of *Xist* (Figure 5C). Notably, RT-qPCR analysis revealed that chronic morphine treatment induced a *Smchd1* downregulation that persisted upon differentiation to mEpiLCs (Figure 5D).

**Figure 5.**
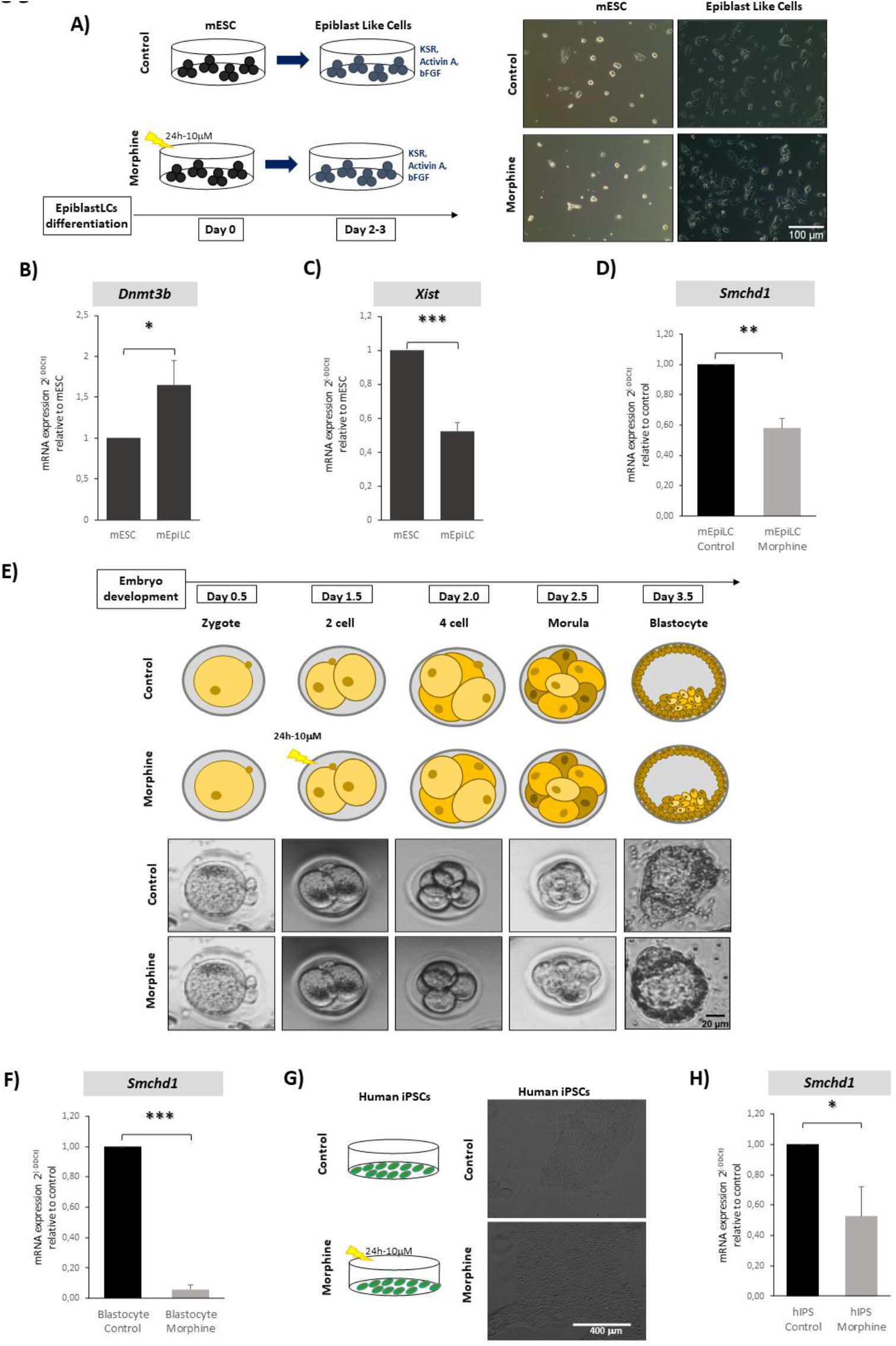
Expression of *Smchd1* across Across Developmental Stages and Species. **(A)** Schematic overview of the differentiation of mESCs into epiblast-like cells (mEpiLCs) following morphine exposure, with representative bright-field images. Scale bar 100 µm. RT-qPCR analysis of **(B)** *Dnmt3b* and **(C)** *Xist* in mESCs and mEpiLCs. **(D)** RT-qPCR analysis of *Smchd1* expression in mEpiLCs derived from control and morphine-exposed mESCs. **(E)** Experimental design for *ex vivo* embryo culture with morphine treatment from the 2-cell stage to blastocyst, with representative images. Scale bar 20 µm. **(F)** RT-qPCR quantification of *Smchd1* expression in individual blastocysts derived from control and morphine-exposed 2-cell embryos. **(G)** Schematic of morphine treatment in human induced pluripotent stem cells (hiPSCs), with representative colony images. Scale bar 400 µm. **(H)** RT-qPCR analysis of *Smchd1* expression in hiPSCs following morphine exposure. Statistical significance was set at *p < 0.05, **p < 0.01, ***p < 0.001.

To examine whether morphine-induced *Smchd1* downregulation persists during early differentiation, we performed *ex vivo* experiments with mouse embryos. Two-cell stage embryos were exposed to chronic morphine treatment and, after morphine withdrawal, were subsequently monitored throughout preimplantation development up to the blastocyst stage at 3.5 days post-fertilization. Embryos progressed normally through cleavage, compaction, and blastocyst formation, with no overt morphological abnormalities relative to control (Figure 5E). Despite this phenotypic normalcy, RT-qPCR analysis revealed a pronounced reduction in *Smchd1* transcript levels in morphine-exposed blastocysts (Figure 5F), indicating that morphine induces sustained epigenetic repression of *Smchd1*that also persists during early embryonic development.

Finally, we examined whether the repression of *Smchd1* induced by morphine is conserved in humans by analyzing morphine-treated human induced pluripotent stem cells (hiPSCs). As we expected, morphine exposure did not alter colony morphology, or cellular appearance, and treated cells retained the typical undifferentiated phenotype (Figure 5G). Nevertheless, RT-qPCR analysis revealed a reproducible decrease in *Smchd1* expression in morphine-treated hiPSCs (Figure 5H), indicating that morphine induces a conserved repression of *Smchd*1 across both developmental stages and species.

## 3. DISCUSSION

Cellular memory is a fundamental property of embryonic development, allowing cells to preserve lineage-specific transcriptional programs long after the initiating signals have disappeared^3^. This persistence is ensured by epigenetic mechanisms—such as DNA methylation, histone modifications, and higher-order chromatin organization—that are mitotically heritable and capable of stabilizing transcriptional states across cell divisions^37^. While cellular memory has traditionally been studied in the context of developmental cues, growing evidence suggests that environmental exposures can also engage epigenetic mechanisms to generate long-lasting molecular states that can impact on cellular phenotypes and therefore causes different diseases^38^. However, the identity of chromatin regulators that translate transient environmental stimuli into durable epigenetic memory remains poorly understood. Transcriptional dynamics suggest a biphasic response to morphine, indicative of transcriptomic reprogramming consistent with a potential long-term epigenetic cellular memory. Morphine elicits long-lasting deregulation of multiple chromatin and cell fate regulators, including neuronal determinants. These results are in line with the observation that prenatal morphine exposure reduces the developmental mass of major organs, including the brain, kidneys, and liver in rodent models, and more critically disrupts normal neuron development, leading to delayed maturation of the nervous system^39^.

Morphine also deregulates transcriptional and epigenetic modifiers, implying a disruption of epigenetic mechanisms that may underlie cellular memory. Specifically, we identify SMCHD1 as a central epigenetic target of morphine-induced epigenetic memory. SMCHD1 (structural maintenance of chromosomes flexible hinge domain containing 1) is a non-canonical SMC family protein that functions as a higher-order chromatin regulator essential for X-chromosome inactivation (XCI), genomic imprinting, and repression of select autosomal gene clusters^40^. Our findings demonstrate that morphine exposure leads to a sustained repression of *Smchd1* expression, persisting for multiple cell divisions after drug withdrawal, thereby fulfilling a key criterion of epigenetic memory. Chronic morphine exposure, therefore, can relieve repression of SMCHD1-regulated gene clusters, leading to transcriptional reprograming that can persist even in the absent of drug. Mechanistically, our data indicate that morphine represses *Smchd1* through chromatin-based mechanisms, prominently involving H3K27me3 enrichment and altered DNA methylation at the *Smchd1* locus. Increased the levels of H3K27me3 repressive histone mark at the *Smchd1* promoter suggests Polycomb-mediated repression as an early response to morphine exposure. Importantly, this repression is not transient, as reduced *Smchd1* expression is retained for at least ten cell cycles after morphine withdrawal, indicating that morphine induces a heritable chromatin state rather than an acute transcriptional response. The enhanced sensitivity of *Smchd1* haploinsufficient mESCs further supports the notion that *Smchd1* dosage is a critical vulnerability point through which environmental exposures can exert long-term epigenetic effects.

The functional consequences of SMCHD1 repression are particularly pronounced in the context of morphine exposure, which specifically reduces DNA methylation at CTCF-bound regions within *Smchd1* target loci. SMCHD1 acts downstream of primary epigenetic marks—such as DNA methylation and H3K27me3—to stabilize transcriptional silencing through higher-order chromatin compaction and restriction of CTCF-mediated chromatin interactions^40,41^. Hypomethylation at these CTCF sites compromises a key recruitment and stabilization signal for SMCHD1, thereby weakening its capacity to antagonize CTCF and maintain repressive 3D genome architecture. Genome-wide studies indicate that SMCHD1 preferentially binds regulatory regions overlapping or adjacent to CTCF sites, where it normally restricts cohesin-driven loop extrusion and long-range enhancer–promoter interactions^40^. Loss of SMCHD1 has been associated with increased CTCF occupancy and altered chromatin architecture at these loci, consistent with roles in maintaining repressive chromatin domains that limit CTCF/cohesin function^41^. Morphine-induced hypomethylation specifically at the CTCF-containing regulatory region of the *Smchd1* gene could therefore reduce the recruitment and stable association of SMCHD1, compounding the effects of direct SMCHD1 repression by the drug. This dual perturbation—diminished SMCHD1 levels and hypomethylation at its CTCF regulatory sites— would be expected to increase CTCF occupancy, facilitate the formation of new chromatin loops, and destabilize inactive compartments, driving them toward active states^40,41^. These findings suggest that morphine-induced repression of SMCHD1 disrupts higher-order chromatin compaction and cluster-level gene silencing, thereby converting a transient environmental stimulus into persistent transcriptional signatures.

Chronic morphine exposure disrupts chromatin regulation of XCI, highlighting the susceptibility of epigenetic memory mechanisms to environmentally induced perturbations. XCI is a paradigmatic example of epigenetic memory in mammals, ensuring stable, heritable silencing of one X chromosome to achieve dosage compensation across development and mitoses^42,43^. This process is established early in embryogenesis and is maintained by a cascade of chromatin modifications and higher-order architectural factors that lock in a silent state for the inactive X (Xi)^42,44^. Although initiation of XCI driven by *Xist* expression is largely unaffected after morphine exposure, it selectively alters the maintenance phase of the process, where SMCHD1 is indispensable for the maintenance of silencing^45^. Together with *Smchd1* repression, morphine causes a destabilization of heterochromatic features characteristic of the Xi, inducing global DNA hypomethylation and a reduced H3K27me3 deposition along the whole X chromosome Moreover, morphine-induced hypomethylation at CTCF-containing regulatory regions of the gene compromises higher-order chromatin organisation for X-linked gene silencing, as SMCHD1 is required to repress CTCF binding and collapse canonical topologically associating domains (TADs)—chromatin regions in which sequences interact more frequently with each other than with neighbouring domains—on the Xi^44^. Consequently, morphine may perturb the architectural reinforcement of XCI, by changing chromatin marks associated with X-linked gene silencing^40,44^.

Beyond XCI, morphine also impacts genomic imprinting, another example of epigenetic memory which involves parent-of-origin monoallelic expression of genes established during gametogenesis^13^. Our results demonstrate the existence of a morphine-induced epigenetic memory through imprinting mechanisms. Chronic morphine exposure induces long-lasting, cluster-specific transcriptional and chromatin changes in imprinted genes, with the *Snrpn* cluster exhibiting unique and persistent dysregulation that induces temporally distinct alterations at both maternally and paternally expressed genes belonging to cluster. These findings suggest for the first time the dynamic and gene-specific responsiveness of imprinted loci to environmental exposure, reflecting the capacity of imprinting to act as a temporal and memory-like regulatory system sensitive to morphine exposure during development. Epigenetic profiling revealed that H3K27me3 marks, rather than DNA methylation alterations at ICR could explain the cluster morphine-induced transcriptional deregulation of *Snrpn* cluster, as imprinted genes are often organized in clusters regulated by ICRs, where DNA methylation and histone modifications such as H3K27me3 and H3K4me2/3 ensure proper allele-specific expression^46^. However, studies in mouse oocytes have demonstrated that allelic identity at the *Snrpn* locus can be recognized prior to the acquisition of DNA methylation, indicating that epigenetic memory is initially stored in a methylation-independent form. At *Snrpn* cluster, DNA methylation appears to function mainly as a stabilizing mechanism rather than the initial carrier of epigenetic information^47^. This indicates histone-based mechanisms independent of DNA methylation may be important at the first phase of the imprinting process after morphine exposure.

Accumulating evidence indicates that chromatin-based mechanisms can mediate DNA methylation–independent imprinting, establishing noncanonical imprinting in early embryos and extraembryonic tissues and contributing to epigenetic memory beyond DNA methylation^48,49^. In this context, long non-coding RNAs have been shown to recruit chromatin modifiers involved in imprinting regulation, including SMCHD1, a critical regulator of the *Snrpn* imprinted cluster ^50,51^. Genetic studies in mice have demonstrated that loss of *Smchd1* results in loss of imprinting (LOI) at the *Snrpn* locus without detectable changes in differential DNA methylation at the primary ICR, proving that *Smchd1* enforces imprinting through mechanisms independent of canonical DNA methylation^52,53^. Consistent with these observations, chronic morphine treatment compromises imprinting at the *Snrpn* cluster through morphine-induced repression of *Smchd1*. Mechanistically, SMCHD1 binds at or near regulatory elements within imprinted clusters and interacts with higher-order chromatin architectural factors, where it is proposed to antagonize CTCF-mediated looping to reinforce a repressive chromatin state^40^. Morphine can destabilize *Smchd1*-mediated chromatin compaction required for the stable maintenance of imprinting at this locus, as we observe morphine-associated hypomethylation at CTCF-containing regulatory regions. *Smchd1* deficiency, together with alterations in CTCF regulatory region, likely represent an early and critical phase of imprinting destabilization at the *Snrpn* cluster after morphine exposure. Remarkably, such perturbations may persist over time through sustained morphine-induced repression of *Smchd1*. These findings support a model in which morphine disrupts genomic imprinting by interfering with *Smchd1*-mediated chromatin repression. This methylation-independent component of epigenetic memory may subsequently influence the maintenance or establishment of DNA methylation, supporting a hierarchical model in which chromatin organization and histone modifications act as primary carriers of imprinting information and revealing a mechanistic link between environmental exposure and the perturbation of imprinting-dependent epigenetic memory.

In summary, our study demonstrates that morphine exposure induces a durable epigenetic memory by targeting higher-order chromatin regulatory mechanisms rather than transient transcriptional responses. We identify sustained repression of *Smchd1* as a key molecular event that converts a short-lived environmental stimulus into long-lasting transcriptional and architectural reprogramming, affecting both X-chromosome inactivation and genomic imprinting. By destabilizing SMCHD1-dependent chromatin compaction, morphine disrupts the maintenance of repressive chromatin domains at CTCF-regulated loci, leading to persistent transcriptional deregulation of gene clusters, including the *Snrpn* imprinted domain. These findings establish a mechanistic link between environmental exposure, chromatin architecture, and epigenetic memory, highlighting how perturbation of architectural epigenetic regulators can translate environmental signals into long-term developmental and transcriptional consequences.

## 4. METHODS

### 4.1 mESC culture and morphine exposure

mESCs (female Oct4-GFP; PCEMM08, PrimCells) were maintained under feeder-free conditions using dishes pre-coated with 0.1% gelatin (G189, Sigma-Aldrich). The cells were cultured in KnockOut DMEM (Gibco, 10829-1018) enriched with 15% KnockOut Serum Replacement (KSR; Gibco, 10828-028), and supplemented with 1% sodium pyruvate (S8636), 1% non-essential amino acids (M7145), 1% penicillin–streptomycin (P4333), 1% L-glutamine (G7513), and 0.07% β-mercaptoethanol (146250), all from Sigma-Aldrich. For maintenance of pluripotency, medium was additionally supplemented with the 2i + LIF cocktail: 1000 U/mL leukemia inhibitory factor (ESG1107), 10 μM PD0325901 (Stemgent, 04-0006), and 30 μM CHIR99021 (StemCell, BX33140). Routine passaging was performed every 48 hours using TrypLE Express Enzyme (ThermoFisher, 12605028) for enzymatic detachment. Oct4-GFP expression was monitored to ensure preservation of the undifferentiated state. For chronic morphine treatment, cells were cultured for 24 hours in medium containing either 10 µM morphine sulfate (Alcaliber) or 0.9% (p/v) NaCl as vehicle control, based on the protocol from Yang et al. (2019)^30^. Following exposure, cells were harvested immediately (P1) or washed and maintained in standard culture conditions for an additional 48 h (P2) or 96 h (P3) before further analysis.

### 4.2. SmcHD1 Knockout of mESCs via CRISPR-Cas9

CRISPR-Cas9-mediated knockout (KO) strategy was implemented on mESCs, based on the protocol described by Gdula et al. (2019)^33^. Three single guide RNAs (sgRNAs) targeting the MommeD1 mutation (*Smchd1*) were designed and cloned into the eSpCas9-2A-puro plasmid (PX459 V2.0, 9.2 kb) using the GenScript services (Supplementary Data 4A-B). The sgRNA sequences used were: caccgTTACAGTGGACGATCAGT, caccgCTGCAATCCACAGTATAGAC, and caccGGCGATCCAGATCAACATATATC. Transfection was performed in the OCT4-GFP mESC line using Lipofectamine 3000 (Thermo Fisher Scientific) according to the manufacturer’s protocol. The transfection mixture was prepared by separately preparing two tubes per well, each containing 50 µl of Opti-MEM (Gibco): one tube with 3 µl of Lipofectamine 3000 reagent and the other with 2 µl of P3000 reagent and 1 µg of PX459 plasmid DNA. The contents of the two tubes were then combined, gently mixed, and incubated for 5–10 minutes. The resulting mixture was added dropwise to pre-seeded cells (20–30% confluence) cultured in 24-well plates coated with 0.1% gelatin. The culture medium was the same as that used for mESC culture. Following 24-48 hours incubation at 37 °C with 5% CO_2_, transfected cells were selected using 4 μg/ml puromycin. After approximately 3 days of selection, puromycin-resistant colonies were expanded, and total RNA was extracted for RT-qPCR analysis to confirm *Smchd1* gene silencing.

### 4.3. mESCs morphine exposure and their differentiation into EpiLCs

mESCs were differentiated into epiblast-like cells (EpiLCs) using the protocol described by Hayashi and Saitou (2013)^54^, to assess the effects of morphine exposure during differentiation. Two experimental groups were established: untreated control mESCs and mESCs exposed to morphine. After chronic morphine treatment for 24h and prior to cell differentiation, 12-well plates (3.5 cm^2^/well) were coated with human plasma fibronectin (16.6 μg/ml in 1× PBS). Approximately 1×10^5^ mESCs were seeded per well in N2B27-based differentiation medium. N2B27 was prepared by mixing equal volumes of two basal media: (1) DMEM/F12+N2, consisting of 494 ml DMEM/F12, 5 ml N2 supplement, 0.5 ml insulin (25 mg/ml), 0.5 ml apo-transferrin (100 mg/ml), 0.33 ml BSA (7.5%), 16.5 μl progesterone (0.6 mg/ml), 50 μl putrescine (160 mg/ml), and 5 μl sodium selenite (3 mM); and (2) Neurobasal+B27, consisting of 480 ml Neurobasal medium, 10 ml B27 supplement, 5 ml penicillin/streptomycin (100 U/ml – 100 μg/ml), and 5 ml L-glutamine (200 mM). The final differentiation medium was supplemented with 1% KnockOut Serum Replacement (KSR), 12 ng/ml basic fibroblast growth factor (bFGF), 20 ng/ml Activin A, and 5 mM β-mercaptoethanol. Cultures were maintained at 37 °C in a humidified incubator with 5% CO_2_, and medium was replaced daily for 2–3 days. Cells from both experimental conditions and differentiation state were collected, processed and stored at −80 °C for further analyses.

### 4.4. Mouse Embryo collection, culture and morphine exposure

Female Swiss Webster (SW) mice (8-10 weeks old) were housed in groups of ten under standard conditions (approx. 23 °C, 12-hour light/dark cycle, lights on 8:00-20:00), with ad libitum access to food and water. All animal procedures were adhered to institutional guidelines and approved by the Animal Ethics Committee of the University of the Basque Country (MA20/2016/142). Superovulation was induced via intraperitoneal injections of 5 IU pregnant mare serum gonadotropin (PMSG; G4527, Sigma-Aldrich), followed 48 hours later by 5 IU human chorionic gonadotropin (hCG; C8554, Sigma-Aldrich). Females were immediately paired with fertile males overnight and successful mating was confirmed by the presence of vaginal plugs the following morning. Approximately 12 hours post-fertilization, mated females were euthanized by cervical dislocation, and zygotes were collected from the oviduct ampullae and placed in EmbryoMax KSOM medium (MR-020P-5F, Merck-Millipore). Cumulus cells were removed using hyaluronidase (H1136, Sigma-Aldrich), and embryos were rinsed twice in fresh KSOM medium and cultured in 40 µL KSOM drops overlaid with 3 mL paraffin oil (18512, Sigma-Aldrich) in culture-grade dishes (430166, Corning). Two-cell-stage embryos were randomly divided into two groups: one untreated group and the other exposed to chronic morphine treatment. After, embryos were washed and transferred to fresh KSOM drops, where they were cultured until reaching the blastocyst stage in a humidified incubator (37 °C, 5% CO_2_) (MCO-15AC, Sanyo). Embryonic development was assessed every 12 hours, recording progression through the 2-cell, 4-cell, morula, and blastocyst stages. Upon reaching the blastocyst stage, embryos were collected and stored at −80 °C for further analyses.

### 4.5. Human Induced Pluripotent Stem Cells (hiPSCs) culture and morphine exposure

Human iPSCs line CREM003i-BU3C2 ^55^, originating from a blood sample of a 40-year-old male was cultured at 36 °C and 0.5% CO_2_. The cells were maintained on Matrigel-coated plates (hESC-qualified, 354277, Corning) using mTeSR1 medium (85870, StemCell Technologies) according to the manufacturer’s protocol. Culture medium was refreshed daily to support optimal growth and pluripotency. Cell passaging was carried out every 5 to 7 days using ReLeSR (05872, StemCell Technologies), a gentle, enzyme-free dissociation method. For chronic morphine exposure experiments, hiPSCs were cultured in standard conditions with the addition of 10 µM morphine sulfate (Alcaliber) for 24 hours. Control cells received a corresponding volume of 0.9% NaCl (p/v) (31414, Sigma-Aldrich) as vehicle. Treatment conditions followed those previously described by Yang et al. (2019)^30^. After the 24-hour exposure period, cells from both groups were collected, processed and stored at −20 °C for RT-qPCR analyses.

### 4.6. RNA-Seq, ChIP-Seq and WGBS data

RNA-Seq, H3K27me3 ChIP-Seq and WGBS data profiles were obtained from National Center of Biotechnology Information (NCBI), Gene Expression Omnibus (GEO) database (https://www.ncbi.nlm.nih.gov/geo/query/acc.cgi?acc=GSE151234 ; Muñoa-Hoyos et al. 2020^31^ and https://www.ncbi.nlm.nih.gov/geo/query/acc.cgi?acc=GSE292082 ; Araolaza et al. 2025^56^. The profile comprised, a poly(a) RNA-Seq experiment, a H3K27me3 repressive histone mark ChIP-Seq experiment and a global methylation WGBS analysis with 4 mESCs samples each, of which 2 were control samples and 2 were morphine treated samples. Both experiments were based on the Illumina Hi-Seq 2500 sequencer with a minimum of 50 million reads per replicate. For data processing, raw data underwent several steps such as quality assessment, UCSC mm10 reference genome alignment, normalization, and identification, as previously described^31,56^. The selection criteria for significant differentially expressed genes (DEGs), differentially binding sites (DBSs) and differentially methylated cytosines (DMC) and genes (DMG) between the control and morphine treated samples were p-value ≤ 0.05 and false discovery rate (FDR) value ≤ 0.05. Finally, an integrative analysis for both sequencing techniques was performed using Venny tool and The Gene Ontology Resource from the GO Consortium (https://geneontology.org/) was used to identify the biological functions.

### 2.7. RNA extraction, quality control and Real-time PCR (RT-qPCR)

Total RNA was isolated from mESCs, EpiLCs, and hiPSCs using TRIzol reagent (15596026, Thermo Fisher Scientific), following the manufacturer’s instructions. Additionally, RNA from blastocyst-stage embryos was specifically extracted with the Arcturus PicoPure RNA Isolation Kit (KIT0103, Thermo Fisher), according to the provided protocol. RNA concentration was measured by absorbance at 260 nm, and purity was verified by calculating the 260/280 nm ratio. cDNA synthesis was reverse transcribed using the iScript cDNA Synthesis Kit (1708891, Bio-Rad) for mESCs, EpiLCs and hiPSCs samples. For blastocyst-derived RNA, cDNA was generated with the SMARTer Universal Low Input RNA Kit (Cat. # 634940, TaKaRa), ensuring compatibility with low-input material. Quantitative real-time PCR (qPCR) was conducted using iTaq Universal SYBR Green SuperMix (1725121, Bio-Rad) on an ABI Prism 7000 Sequence Detection System. The thermal cycling program included an initial denaturation at 95 °C for 10 minutes, followed by 39 cycles of 95º for 20 seconds (denaturation) and 59 °C for 1 minute (annealing and extension). All reactions were performed in triplicate and repeated across at least three independent biological replicates. Relative gene expression was calculated using the ΔΔCt method. For normalization *Gapdh* and *Pcx* were employed as internal reference genes in mouse samples, while beta-actin (*Actb*) and hypoxanthine phosphoribosyltransferase 1 (*Hprt1*) were used for human-derived samples. The complete list of primer sequences is provided in Supplementary Data 6.

### 4.8. Protein extraction, quality control and Western Blot

For protein analysis, whole-cell extracts were prepared from approximately 50,000 mESCs per sample. Extracts were mixed with 4× loading buffer containing 10% (v/v) dithiothreitol (DTT) and denatured by boiling at 95 °C for 10 minutes. Proteins were then separated by one-dimensional SDS-PAGE using 12% resolving gels and transferred onto PVDF membranes (Amersham Hybond, Sigma) using the Mini Trans-Blot system (Bio-Rad Laboratories, Hercules, CA, USA). Membranes were blocked for 1 hour in Blotto solution (20 mM Tris–HCl pH 7.5, 0.15 M NaCl, 1% Triton X-100) supplemented with 5% bovine serum albumin (BSA) to prevent nonspecific binding. Primary antibody incubation was performed overnight at 4 °C using a monoclonal rabbit anti-H2AK119ub antibody (D27C4, Cell Signaling) and a polyclonal rabbit anti-H3K27me3 antibody (A2363, Abclonal), both diluted 1:1000. To verify equal protein loading and normalize signal intensity, membranes were reproved with a monoclonal mouse anti-beta actin peroxidase antibody (A3854, Sigma) at a 1:25,000 dilution and a polyclonal rabbit anti-H3 antibody (Ab1791, Abcam) at a dilution 1:5000. After three washes (5 minutes each) in Blotto, membranes were incubated for 1 hour at room temperature with an HRP-conjugated goat anti-rabbit secondary antibody (Santa Cruz Biotechnology, sc-2004) diluted 1:1000 in Blotto containing 5% BSA. Protein bands were visualized using enhanced chemiluminescence on a ChemiDoc XRS imaging system (Bio-Rad).

### 4.9. Sequencing data statistical analysis

For all sequencing modalities, read quality and adapter trimming were performed using FastQC (v0.11.6^57^) and Trim Galore! (v0.4.5/v0.6.2^58^). Filtered reads were mapped to the mouse genome (mm10) using technique-specific aligners as described below.

#### RNA-Seq

Mapping was executed with HISAT2 (v2.1.0^59,60^). Gene annotation was performed against the UCSC table browser using featureCounts (v1.16.0^61^). DGE analysis was assessed via edgeR (v3.20^62^) using a quasi-likelihood framework and a negative binomial distribution to account for biological over-dispersion. DEGs with an adjusted p-value < 0.05 were defined as significant.

#### ChIP-Seq

Trimmed reads were aligned using the Bowtie2 software (v2.3.4^63^). Peak calling was performed using MACS (v1.4.1^64^), based on a Poisson distribution model. Following peak calling, DBSs between control and morphine-treated samples were identified using DiffBind package (v2.8.0^65,66^), applying a negative binomial distribution to model normalized read counts within peak regions to account for biological variability. DBSs with an adjusted p-value < 0.05 were defined as significant. Finally, peaks were annotated to the nearest gene and TSS via ChIPSeeker (v1.16.1^67^), considering an FDR < 0.05 as the threshold for significance.

#### WGBS

Pre-processed reads were mapped, deduplicated and methylation extracted using Bismark (v0.22.1^68^). Methylation counts were obtained with methylDackel (v0.5.1; https://github.com/dpryan79/MethylDackel) and DMCs were identified through a consensus approach using two strategies: 1) a logistic regression model in methylKit (v1.16.1^69^) and 2) a binomial distribution in edgeR (v3.32.1^62,70^). DMCs with an adjusted p-value < 0.05 were defined as significant. Only DMCs identified by both tools (via Venny^71^) meeting the thresholds of 25% methylation difference with methylKit and log2 fold-change ≥2 with edgeR, were retained for downstream analysis.

P-values from all -omics were adjusted for multiple testing using the Benjamini-Hochberg procedure to evaluate FDR, considering an FDR<0.05 as the threshold for significance. Downstream, the biological functional analysis was conducted through The Gene Ontology Resource from the GO Consortium (GO v16.1.0; http://geneontology.org)^72^. Imprinted genes were categorized in maternally or paternally expressed genes, using the Geneimprint^73^ and Catalogue of Parent of Origin Effects databases (http:(//igc.otago.ac.nz).

Statistical differences between the control and experimental groups for qPCR analyses were analyzed using a two-tailed unpaired Student’s t-test, from at least five independent experiments. P-values < 0.05 were considered statistically significant.

## ACKNOWLEDGEMENTS

The authors particularly acknowledge SGIKer resources of UPV/EHU for technical support with the computational calculations, which were carried out in the Arina informatics cluster. During the preparation of this manuscript/study, the authors used IA (OpenAI, GPT-4o) for the purposes of improving the English writing. The authors have reviewed and edited the output and take full responsibility for the content of this publication.

## FUNDING INFORMATION

This work was supported by MCIU, Instituto de Salud Carlos III and co-funded by European Union (PI24/01131) to NS and Basque Government, Department of Education (IT1966-26) to NS, IMH and MA. The work was also supported by Researcher Fellowships from Basque Government MA (Mikel Albizuri) and from UPV/EHU to IC.

## AUTHOR CONTRIBUTIONS

Conceptualization, I.M.-H., M.A. (Manu Araolaza) and N.S.; methodology, I.M.-H., M.A. (Manu Araolaza), P.G. and N.S.; software, I.M.-H.; formal analysis, I.M.-H. and M.A. (Manu Araolaza); investigation, I.M.-H., M.A. (Manu Araolaza), I.C. and M.A. (Mikel Albizuri); resources, N.S.; data curation, I.M.-H.; writing—original draft preparation, I.M.-H. and M.A. (Manu Araolaza); writing—review and editing, I.M.-H., M.A. (Manu Araolaza), P.G. and N.S.; visualization, I.M.- H., M.A. (Manu Araolaza), I.C., M.A. (Mikel Albizuri) and P.G.; supervision, N.S. and P.G.; project administration, N.S.; funding acquisition, N.S. All authors have read and agreed to the published version of the manuscript.

## COMPETING INTERESTS

The authors declare no conflicts of interest.

## Notes

### Competing Interest Statement

The authors have declared no competing interest.

