## Supplementary Information for "“Chronic morphine treatment induces a conserved Smchd1-dependent epigenetic memory that disrupts X-chromosome inactivation and genomic imprinting”"

### ABSTRACT

Epigenetic memory ensures stable inheritance of gene expression patterns critical for embryonic development. Environmental exposures can disrupt this memory, yet the mechanisms remain unclear. Here we demonstrate that chronic morphine exposure induces a persistent transcriptomic and epigenetic memory by repressing *Smchd1*, a key chromatin regulator, in mouse embryonic stem cells, preimplantation embryos, and human induced pluripotent stem cells. This repression compromises maintenance of X-chromosome inactivation and genomic imprinting, leading to sustained dysregulation of developmentally important gene clusters. Morphine-induced epigenetic alterations also involve changes in DNA methylation and histone modifications along the X chromosome and notably increased H3K27me3 at the *Smchd1* locus. These findings reveal a conserved mechanism by which opioid exposure disrupts higher-order chromatin architecture and epigenetic memory during early development, potentially contributing to long-term developmental and clinical outcomes.

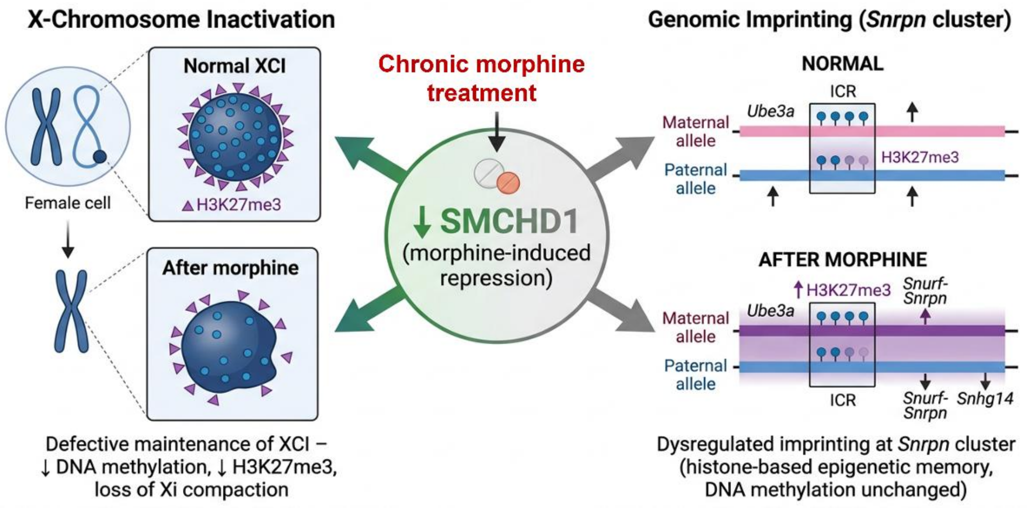

### Supplementary Data 1

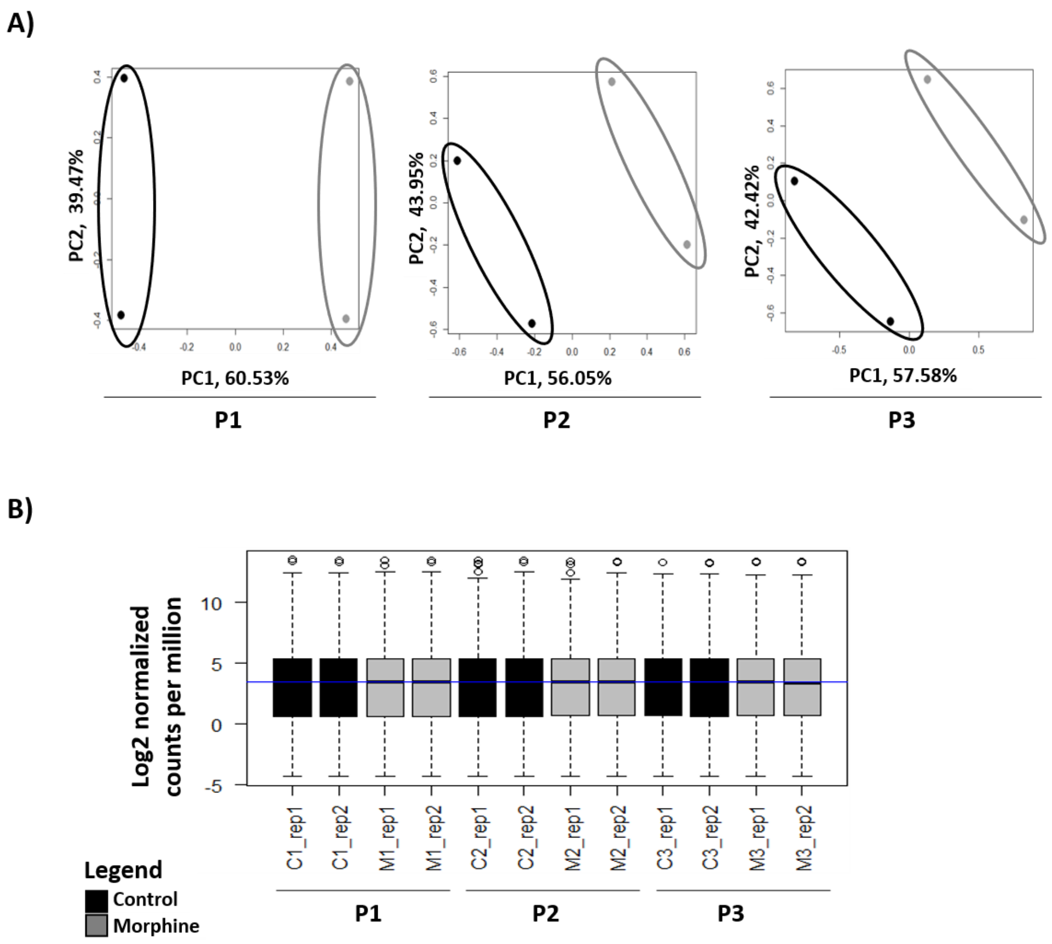


**Supplementary Figure 1. RNA-seq quality control** obtained from datasets at passages P1, P2, and P3 following chronic morphine exposure in mESCs. (**A**) Principal component analysis (PCA) plots showing sample clustering for control and morphine-treated conditions at each passage, with the percentage of variance for each principal component indicated. (**B**) Boxplots displaying $log_2$ normalized counts per million reads (CPM) for all samples after TMM normalization, shown separately for each passage and condition.

### Supplementary Data 2

**Supplementary Table 1. Persistent DEGs induced by morphine across the time identified by hierarchical clustering.** Table listing the 181 DEGs grouped according to clusters which shares expression patterns over time. Cluster 1:  upregulated genes immediately following morphine exposure but progressively downregulated over the time. Cluster 2: downregulated genes that increased their expression over the time. For each gene, the full gene name and molecular function, derived from Gene Ontology annotations are provided.

| Gene | Complete gene name | Molecular Function (GO:MF) |
|---|---|---|
| **Cluster 1** | | |
| ***Aprt*** | Adenine phosphoribosyltransferase | transferase activity; protein binding; catalytic activity |
| ***Atp5d*** | ATP synthase F1 subunit delta | ATP binding; structural constituent of mitochondrial ATP synthase; protein binding |
| ***Bola2*** | BolA family member 2 | protein binding; metal ion binding; chaperone binding |
| ***Chpf*** | Chondroitin polymerizing factor | glycosyltransferase activity; protein binding; transferase activity |
| ***Ccdc124*** | Coiled-coil domain containing 124 | protein binding; microtubule binding; coiled-coil binding |
| ***Dohh*** | Deoxyhypusine hydroxylase | oxidoreductase activity; protein binding; catalytic activity |
| ***Elob*** | Elongin B | protein binding; transcription cofactor activity; ubiquitin ligase regulator activity |
| ***Eif1*** | Eukaryotic translation initiation factor 1 | RNA binding; translation initiation factor activity; protein binding |
| ***Eif4ebp1*** | Eukaryotic translation initiation factor 4E binding protein 1 | protein binding; translation regulator activity; RNA binding |
| ***Epop*** | Elongin BC-interacting protein | chromatin binding; transcription cofactor activity; protein binding |
| ***Fasn*** | Fatty acid synthase | transferase activity; acyltransferase activity; protein binding |
| ***Hspb1*** | Heat shock protein family B (small) member 1 | protein binding; unfolded protein binding; chaperone binding |
| ***Jund*** | Jun D proto-oncogene | DNA-binding transcription factor activity; transcription coregulator activity; protein binding |
| ***Mrpl34*** | Mitochondrial ribosomal protein L34 | RNA binding; structural constituent of ribosome; protein binding |
| ***Nr1h2*** | Nuclear receptor subfamily 1 group H member 2 | DNA-binding transcription factor activity; ligand-activated receptor activity; protein binding |
| ***Pmm1*** | Phosphomannomutase 1 | transferase activity; isomerase activity; protein binding |
| ***Pou5f1*** | POU class 5 homeobox 1 (Oct4) | DNA-binding transcription factor activity; transcription coregulator activity; RNA polymerase II regulatory region sequence-specific binding |
| ***Pycr2*** | Pyrroline-5-carboxylate reductase 2 | oxidoreductase activity; protein binding; NAD(P)H binding |
| ***Rbm38*** | RNA-binding motif protein 38 | RNA binding; protein binding; translation regulator activity |
| ***Rpl11*** | Ribosomal protein L11 | RNA binding; structural constituent of ribosome; protein binding |
| ***Rpl37*** | Ribosomal protein L37 | RNA binding; structural constituent of ribosome; protein binding |
| ***Rpl41*** | Ribosomal protein L41 | RNA binding; structural constituent of ribosome; protein binding |
| ***Rps19*** | Ribosomal protein S19 | RNA binding; structural constituent of ribosome; protein binding |
| ***Slc25a39*** | Solute carrier family 25 member 39 | transporter activity; protein binding; ion binding |

| Gene | Complete gene name | Molecular Function (GO:MF) |
|---|---|---|
| **Cluster 2** | | |
| ***Tcf3*** | Transcription factor 3 | DNA-binding transcription factor activity; transcription coregulator activity; protein binding |
| ***Alms1*** | Alström syndrome 1 | protein binding; centrosome binding; microtubule binding |
| ***Ascc3*** | Activating signal cointegrator 1 complex subunit 3 | DNA binding; helicase activity; protein binding |
| ***Atm*** | ATM serine/threonine kinase | protein kinase activity; DNA binding; protein binding |
| ***Atr*** | ATR serine/threonine kinase | protein kinase activity; DNA binding; protein binding |
| ***Atrx*** | ATRX chromatin remodeler | chromatin binding; ATPase activity; protein binding |
| ***Brca2*** | Breast cancer type 2 susceptibility protein | DNA binding; recombinase cofactor activity; protein binding |
| ***Cbx5*** | Chromobox 5 | chromatin binding; protein binding; transcription corepressor activity |
| ***Cdk2ap2*** | CDK2-associated protein 2 | protein binding; cyclin-dependent kinase inhibitor activity |
| ***Cenpe*** | Centromere protein E | microtubule motor activity; ATP binding; protein binding |
| ***Cenpf*** | Centromere protein F | microtubule binding; protein binding; kinetochore binding |
| ***Cltc*** | Clathrin heavy chain | protein binding; clathrin binding; structural molecule activity |
| ***Csnk2a1*** | Casein kinase 2 alpha 1 | protein kinase activity; ATP binding; protein binding |
| ***Epop*** | Elongin BC-interacting protein | chromatin binding; transcription cofactor activity; protein binding |
| ***Kdm5a*** | Lysine demethylase 5A | histone demethylase activity; protein binding; chromatin binding |
| ***Kdm6a*** | Lysine demethylase 6A | histone demethylase activity; protein binding; chromatin binding |
| ***Kif20b*** | Kinesin family member 20B | microtubule motor activity; ATP binding; protein binding |
| ***Klf2*** | Kruppel-like factor 2 | DNA-binding transcription factor activity; sequence-specific DNA binding; RNA polymerase II transcription regulatory region binding |
| ***Mki67*** | Marker of proliferation Ki-67 | protein binding; nucleolar binding; chromatin binding |
| ***Naa50*** | N-alpha-acetyltransferase 50 | acetyltransferase activity; protein binding; enzyme regulator activity |
| ***Nipbl*** | Nipped-B-like protein | cohesin loading factor activity; DNA binding; protein binding |
| ***Ogt*** | O-linked N-acetylglucosamine transferase | transferase activity; protein binding; catalytic activity |
| ***Pds5a*** | PDS5 cohesin-associated factor A | protein binding; DNA binding; cohesin regulator activity |
| ***Phip*** | Pleckstrin homology domain interacting protein | protein binding; transcription cofactor activity; kinase binding |

| | | |
|---|---|---|
| ***Pola1*** | DNA polymerase alpha 1 | DNA polymerase activity; protein binding; DNA primase activity |
| ***Pou5f1*** | POU class 5 homeobox 1 (Oct4) | DNA-binding transcription factor activity; transcription coregulator activity; RNA polymerase II regulatory region sequence-specific binding |
| ***Psme4*** | Proteasome activator subunit 4 | proteasome binding; protein binding; endopeptidase activator activity |
| ***Rest*** | RE1-silencing transcription factor | DNA-binding transcription factor activity; transcription corepressor activity; protein binding |
| ***Rif1*** | RAP1 interacting factor 1 | protein binding; DNA binding; telomere maintenance factor activity |
| ***Samd1*** | Sterile alpha motif domain containing 1 | protein binding; transcription regulator activity; chromatin binding |
| ***Sf3b1*** | Splicing factor 3b subunit 1 | RNA binding; protein binding; spliceosome component |
| ***Smc3*** | Structural maintenance of chromosomes 3 | ATP binding; protein binding; DNA binding |
| ***Smchd1*** | Structural maintenance of chromosomes flexible hinge domain containing 1 | chromatin binding; ATP binding; protein binding |
| ***Supt16*** | SPT16 homolog | DNA binding; protein binding; chromatin binding |
| ***Suz12*** | SUZ12 polycomb repressive complex 2 subunit | chromatin binding; histone methyltransferase regulator activity; transcription corepressor activity |
| ***Tcf3*** | Transcription factor 3 | DNA-binding transcription factor activity; transcription coregulator activity; protein binding |
| ***Tet1*** | Tet methylcytosine dioxygenase 1 | dioxygenase activity; DNA binding; protein binding |
| ***Top2a*** | DNA topoisomerase II alpha | DNA topoisomerase activity; ATP binding; protein binding |
| ***Tpr*** | Translocated promoter region | nuclear pore complex binding; protein binding; coiled-coil binding |
| ***Trip12*** | Thyroid hormone receptor interactor 12 | ubiquitin ligase activity; protein binding; transcription coregulator activity |
| ***Ube2d3*** | Ubiquitin conjugating enzyme E2 D3 | ubiquitin-protein transferase activity; protein binding; catalytic activity |
| ***Ubn2*** | Ubinuclein 2 | chromatin binding; protein binding; transcription cofactor activity |
| ***Usp9x*** | Ubiquitin specific peptidase 9 X-linked | cysteine-type deubiquitinase activity; protein binding; hydrolase activity |
| ***Wapl*** | WAPL cohesin release factor | protein binding; DNA binding; chromatin binding |
| ***Zfp445*** | Zinc finger protein 445 | DNA-binding transcription factor activity; protein binding; metal ion binding |
| ***Zfp518a*** | Zinc finger protein 518A | DNA-binding transcription factor activity; protein binding; metal ion binding |

### Supplementary Data 3

**Supplementary Table 4. Morphine-sensitive genes at both the transcriptomic and epigenetic levels.** The table lists the 14 genes that overlaps following a multi-omics analysis combining transcriptomic (RNA-seq), methylome (whole-genome bisulfite sequencing, WGBS) and the repressive histone H3K27me3 (ChIP-seq) data. For each gene, the full gene name, molecular function and associated biological processes are provided.

| Gene | Complete gene name | Molecular Function (GO:MF) | Biological Process (GO:BP) |
|---|---|---|---|
| ***Akap11*** | A-kinase anchoring protein 11 | Protein kinase A binding; scaffold protein binding; protein binding | Signal transduction; regulation of protein localization; camp-dependent protein kinase signaling |
| ***Epop*** | Elongin BC-interacting protein (EPOP) | Chromatin binding; protein binding; transcription cofactor activity | Regulation of transcription by RNA polymerase ii; maintenance of pluripotency; histone modification |
| ***Hspb1*** | Heat shock protein family B (small) member 1 | Protein binding; unfolded protein binding; chaperone binding | Response to heat; response to oxidative stress; regulation of apoptosis |
| ***Klf2*** | Kruppel-like factor 2 | DNA-binding transcription factor activity; sequence-specific DNA binding; RNA polymerase II transcription regulatory region binding | Regulation of transcription; blood vessel development; response to shear stress |
| ***Pcm1*** | Pericentriolar material 1 | Centrosome binding; protein binding; microtubule binding | Cell cycle; centrosome organization; ciliogenesis |
| ***Pgap1*** | Post-GPI attachment to proteins 1 | Hydrolase activity; phospholipase activity; protein binding | Gpi anchor metabolic process; protein maturation; membrane organization |
| ***Pikfyve*** | Phosphoinositide kinase, FYVE-type zinc finger containing | Phosphatidylinositol phosphate kinase activity; zinc ion binding; phosphotransferase activity | Endomembrane system organization; endosomal transport; phosphatidylinositol metabolic process |
| ***Pou5f1*** | POU class 5 homeobox 1 (Oct4) | DNA-binding transcription factor activity; transcription coregulator activity; RNA polymerase II regulatory region sequence-specific binding | Cell differentiation; stem cell population maintenance; regulation of transcription involved in cell fate |
| ***Psme4*** | Proteasome activator subunit 4 | Proteasome binding; protein binding; endopeptidase activator activity | Protein catabolic process; dna repair; proteasome-mediated ubiquitin-independent protein degradation |
| ***Smchd1*** | Structural maintenance of chromosomes flexible hinge domain containing 1 | Chromatin binding; ATP binding; protein binding | Gene silencing; X-chromosome inactivation; chromatin organization |
| ***Ssbp4*** | Single-stranded DNA-binding protein 4 | Single-stranded DNA binding; protein binding; transcription factor binding | Embryonic development; regulation of transcription; DNA-templated transcription |
| ***Suz12*** | SUZ12 polycomb repressive complex 2 subunit | Chromatin binding; histone methyltransferase regulator activity; transcription corepressor activity | Histone h3-k27 methylation; gene silencing; embryonic development |
| ***Tead1*** | TEA domain family member 1 | DNA-binding transcription factor activity; cofactor-dependent transcription factor activity; protein binding | Regulation of transcription; muscle organ development; Hippo signaling pathway |
| ***Zfp644*** | Zinc finger protein 644 | DNA-binding transcription factor activity; metal ion binding; protein binding | Regulation of transcription; visual system development; chromatin organization |

### Supplementary Data 4

A)

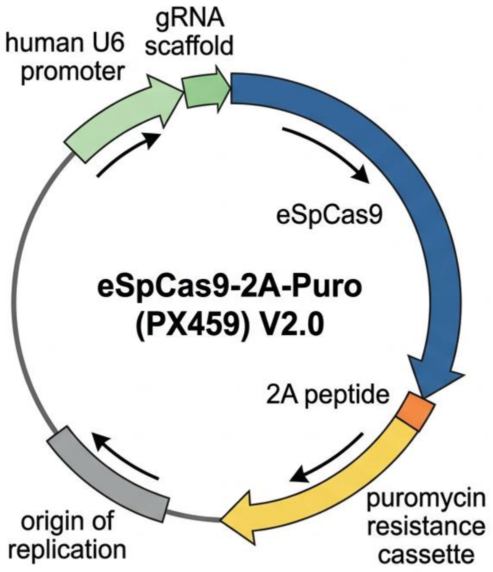


B)

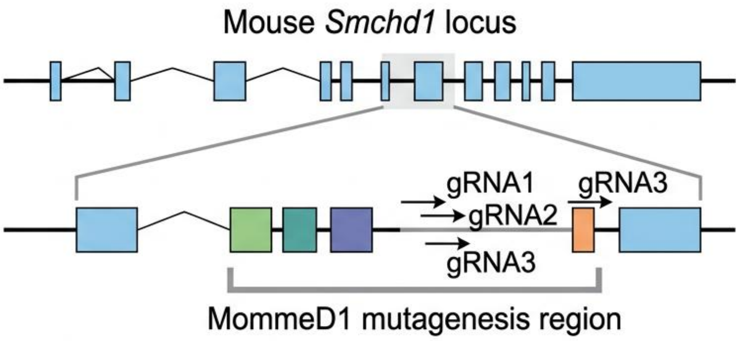


C)

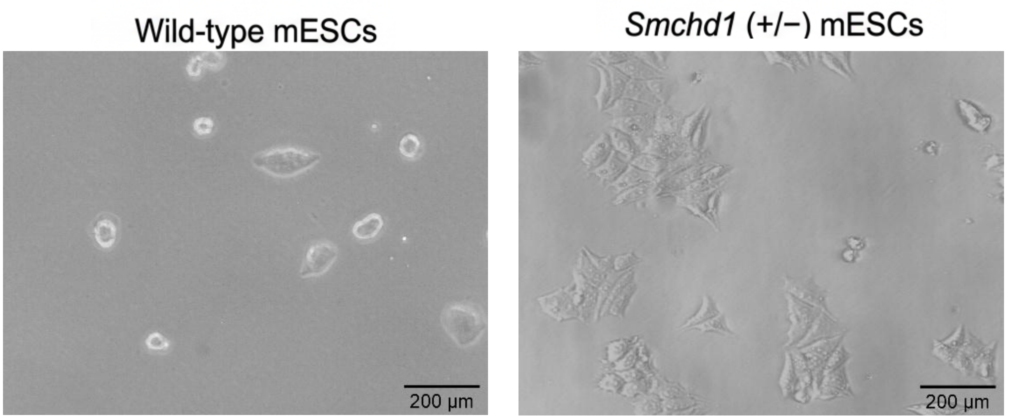


**Supplementary Figure 2. CRISPR/Cas9-mediated targeting of the mouse Smchd1 MommeD1 region in mESCs. (A)** Schematic representation of the eSpCas9-2A-Puro (PX459) V2.0 plasmid used for gene editing, showing the U6 promoter and gRNA scaffold, eSpCas9, 2A peptide and puromycin resistance cassette. **(B)** Linear map of the Smchd1 locus highlighting the MommeD1 region. The three guide RNAs designed to target Smchd1 within the MommeD1 region are indicated by arrows (gRNA1: caccgTTACAGTGGACGATCACGTT; gRNA2: caccgCTGCATCCACCATAGTAGAC; gRNA3: caccGGC GATCCAGATCAAACATC) positioned over the enlarged segment of the gene. **(C)** Representative phase-contrast images of wild-type mESCs and Smchd1 heterozygous knockout [Smchd1 (+/−)] mESCs generated using the PX459 vector and the MommeD1-targeting guide RNAs shown in (A). Scale bar, 200 μm.

### Supplementary Data 5

**A)**

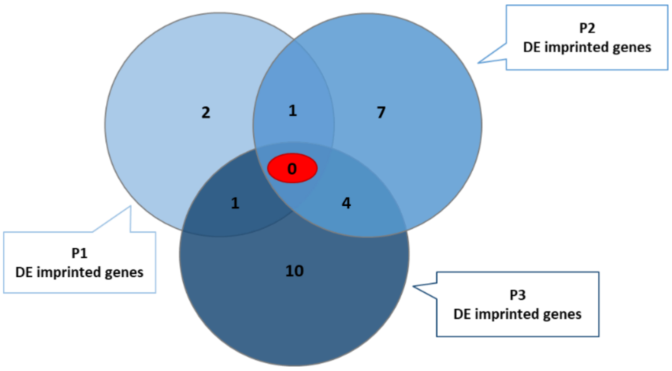


**B)**

| **DE imprinted genes at P1** | | | **DE imprinted genes at P3** | | |
|---|---|---|---|---|---|
| ***Mirg*** | Maternally expressed | Dlk1-Dio3 cluster | ***Meg3*** | Maternally expressed | Dlk1-Dio3 cluster |
| ***Dhcr7*** | Maternally expressed | Kcnq1 cluster | ***Rian*** | Maternally expressed | Dlk1-Dio3 cluster |
| ***Ube3a*** | Maternally expressed | Snrpn cluster | ***Tssc4*** | Maternally expressed | Kcnq1 cluster |
| ***Rhox5*** | Unknowingly expressed | -- | ***Cd81*** | Maternally expressed | Kcnq1 cluster |
| **DE imprinted genes at P2** | | | ***Ube3a*** | Maternally expressed | Snurf-Snrpn cluster |
| ***Snurf*** | Paternally expressed | Snrpn cluster | ***Snhg14*** | Paternally expressed | Snurf-Snrpn cluster |
| ***Snrpn*** | Paternally expressed | Snrpn cluster | ***Peg10*** | Paternally expressed | Sgce-Peg10 cluster |
| ***Snhg14*** | Paternally expressed | Snrpn cluster | ***Zdbf2*** | Paternally expressed | Gpr1-Zdbf1 cluster |
| ***Peg3*** | Paternally expressed | Peg3 cluster | ***Igf2r*** | Maternally expressed | Igf2r-Air cluster |
| ***Ppp1r9a*** | Paternally expressed | Sgce-Peg10 cluster | ***Copg2*** | Maternally expressed | Peg1-Copg2 cluster |
| ***Zdbf2*** | Paternally expressed | Gpr1-Zdbf1 cluster | ***Peg1*** | Paternally expressed | Peg1-Copg2 cluster |
| ***Mcts2*** | Paternally expressed | H13-Mcts2 cluster | ***Tsix*** | Maternally expressed | Tsix-Zcchc13 cluster |
| ***H19*** | Maternally expressed | Igf2-H19 cluster | ***Slc38a4*** | Paternally expressed | -- |
| ***Igf2r*** | Maternally expressed | Igf2r-Air cluster | ***Impact*** | Paternally expressed | -- |
| ***Cobl*** | Maternally expressed | -- | ***Lin28a*** | Unknowingly expressed | -- |
| ***Slc38a4*** | Paternally expressed | -- | | | |
| ***Rhox5*** | Unknowingly expressed | -- | | | |

**Supplementary Figure 3**. **Differentially expressed (DE) imprinted genes at time points P1–P3. (A)** Venn diagram of morphine-induced differentially expressed imprinted genes across three passages (P1-P3). Numbers within each sector indicate the number of genes unique to a given condition (non-overlapping regions) or shared between two or three conditions (overlapping regions). The central red sector highlights the number of DE imprinted genes common to all three conditions. **(B)** List of imprinted genes that are differentially expressed at P1 and P2 and at P3 (bottom), indicating for each gene its parental expression pattern (maternally, paternally or unknowingly expressed) and the imprinted cluster or genomic domain to which it belongs.

### Supplementary Data 6

**Supplementary Table 5. RT-qPCR primer information.** Table containing the primer sequences used for RT-qPCR analyses. Primers were designed for Mus musculus, with *mGapdh* and *mPcx* used as housekeeping genes. For experiments additionally performed in human induced pluripotent stem cells (hiPSCs), *hActb* and *hHprt1* were used as housekeeping controls.

| Gene | Complete name | Sequences |
|---|---|---|
| ***hActb*** | Actin Beta | (f) TGGCATCCACGAAACTACCT |
| | | (r) ACGGAGTACTTGCGCTCAG |
| ***hHprt1*** | Hypoxanthine Phosphoribosyltransferase 1 | (f) CCTGGCGTCGTGATTAGTGA |
| | | (r) CGAGCAAGACGTTCAGTCCT |
| ***hSmcHD1*** | Structural Maintenance Of Chromosomes Flexible Hinge Domain Containing 1 | (f) AGGCAGTGGAGATGTTTTGG |
| | | (r) GCGTCAGTGGTTAGGGTGA |
| ***mDnmt1*** | DNA Methyltransferase 1 | (f) GCCAGTTGTGTGACTTGGAA |
| | | (r) GTCTGCCATTTCTGCTCTCC |
| ***mDnmt3B*** | DNA Methyltransferase 3 Beta | f) ACTTGGTGATTGGTGGAAGC |
| | | (r) CCAGAAGAATGGACGGTTGT |
| ***mEzh2*** | Enhancer Of Zeste 2 Polycomb Repressive Complex 2 Subunit | (f) AGAATGTGGAGTGGAGTGGTG |
| | | (r) CAGTGGGAACAGGTGCTATG |
| ***mGadph*** | Glyceraldehyde 3-phosphate dehydrogenase | (f) TATGACTCCACTCACGGCAAATT |
| | | (r) TCGCTCCTGGAAGATGGTGAT |
| ***mPcx*** | Pyruvate Carboxylase | (f) CAACACCTACGGCTTCCCTA |
| | | (r) CCACAAACAACGCTCCAT |
| ***mSmcHD1*** | Structural Maintenance Of Chromosomes Flexible Hinge Domain Containing 1 | (f) TGGATAGGACAGTAGCCGAGA |
| | | (r) ATTGCTTCCCCCTTTTTGTT |
| ***mXist*** | X Inactive Specific Transcript | (f) GCCTCTGATTTAGCCAGCAC |
| | | (r) GCAACCCCAGCAATAGTCAT |
